# Adipogenic differentiation and inflammatory response is orchestrated by regulating euchromatic histone methyltransferases

**DOI:** 10.1101/2024.06.03.597095

**Authors:** Mahua Chakraborty, Jithu Anirudhan, Vignesh K Krishnamoorthy, Nitya Shree, Shravanti Rampalli

**Affiliations:** Institute for Stem Cell Science and Regenerative Medicine (inStem), Bangalore 560065, India.; Cardiorespiratory Disease Biology, CSIR-Institute of Genomics and Integrative Biology (CSIR-IGIB), New Delhi 110025, India.; Academy of Scientific and Innovative Research (AcSIR), Ghaziabad 201002, India.; School of Regenerative Medicine, Manipal Academy of Higher Education, Bangalore, India.

## Abstract

Euchromatic histone methyltransferases (EHMT1/2) play a key role in adipogenesis by regulating gene expression. While the downstream gene functions of EHMTs in adipogenic differentiation have been studied, their regulation and precise individual contributions remain elusive. We discovered the existence of a regulatory mechanism, wherein EHMT1 governs the interdependent expression of itself and the master regulator PPARƴ during the early phase of adipogenesis. In later stages, EHMT2 levels decline along with reduction in H3K9 dimethylation. Alteration of above sequence of events alone or in the presence of saturated-fatty acids lead to precocious induction of high levels of PPARƴ, accelerated adipogenesis and hypertrophic adipocytes with a pro-inflammatory phenotype. Countering the decrease in EHMTs effectively abrogated the inflammatory response of the adipocytes. Accordingly, induction of obesity by a high fat diet was sufficient to downregulate H3K9me2 levels and expression of EHMTs along with enhanced IL-6 generation. Taken together, our studies reveal a critical regulatory role played by EHMTs, which coordinates adipogenesis and obesity-induced inflammation.

## Introduction

Chromatin modification through histone methylation is one of the major epigenetic mechanisms that alter nucleosome structure to regulate gene expression (1)(2). Euchromatic Histone Methyl Transferases (EHMTs) belong to an evolutionarily conserved family of SET domain-containing proteins, which are involved in methylation of histones (3)(4) and non-histone proteins (eg. p53, Rb, and Hsp90) (5)(6)(7). They play diverse roles in various cellular mechanisms involving cell fate specification, cellular differentiation, tissue development, homeostasis and immune response (8). Therefore, perturbation in their regulation caused by inherited or acquired mutations tips off the balance in the homeostatic mechanism resulting in pathological conditions (9)(10). EHMTI(GLP) is a ubiquitously expressed chromatin modifier, which forms a heteromeric complex with EHMT2(G9a) and brings about mono- or di-methylation of H3K9 of euchromatin. This results in heterochromatin formation and transcriptional repression, thereby regulating the transcriptional output during cellular development and differentiation (11). Although, EHMT1-EHMT2 complex forms are usually found *in vivo* with active methyltransferase activity, EHMT1 also possesses mutually exclusive functions as compared to EHMT2. EHMT1 is critical for B-cell and T-cell, germ cell development (12)(13) and neural development (14)(15) along with brown fat specification (6)(17). Complete deletion of *Ehmt1* leads to embryonic lethality in mice (18), while its haplo-insufficiency is known to cause Kleefstra syndrome (KS) in humans (19). Although the downstream gene functions of *Ehmt1/2* have been extensively studied in pathological conditions, very little is known about the regulation of these chromatin modifiers under homeostatic conditions.

Mounting evidence has implicated the role of post-translational modification of histones by histone methyltransferases as one of the major epigenetic mechanisms functional during adipogenesis (20)(21)(22)(23). Recent studies have established the fact that haploinsufficiency of *Ehmt1* in humans causes obesity and almost 40% of the KS patients are found to be overweight (24). In 2013, using the adipose-specific *Ehmt1* knockout mouse, Ohno et al. had uncovered the role of EHMT1 in BAT mediated thermogenesis. Their study showed that absence of EHMT1 in mouse adipose tissue leads to obesity and systemic insulin resistance (16). In another study Wang et al. had previously shown that EHMT2 represses adipogenesis by downregulating the expression of *Pparf* (*25*). However, the contribution of individual EHMTs and regulation of *Ehmts* in adipogenesis remains unanswered.

In terms of the regulation of *Ehmt1/Ehmt2* function, several studies have claimed the role of conditional binding partners and non-coding RNAs (microRNA). In 2015 two proteins were found - ZNF644 and WIZ, both containing zinc finger motifs that can guide the EHMT1/2 complex to specific sites in the genome to switch off the expression of particular genes during early neurodevelopment (26). Most of the studies dealing with regulation of *Ehmt1/2* have been done in context of some pathological conditions such as Kleefstra syndrome, congenital heart defects, cancers or neurological anomalies. In fact, very little has been investigated related to *Ehmt1/2* regulation under homeostatic conditions such as differentiation during development. In order to address this shortcoming, here we primarily focused on the regulation of histone modifiers *Ehmt1/2* during adipogenic development.

In the current study we provide evidence for the first time, claiming the existence of a new regulatory mechanism for a class of chromatin modifiers, specifically euchromatic histone methyltransferases - EHMT1. We demonstrate that histone methyltransferase *Ehmt1* controls its own expression during early adipogenesis while modulating the expression of the master transcription factor *Pparƴ* via a feedback loop mechanism. During this process, EHMT1 maintains its minimal expression, blunting the repression of critical downstream adipogenic genes, thereby successfully driving adipogenic differentiation. We also demonstrate that the untimely loss of global H3K9me2 activity in preadipocytes induces precocious induction of PPARƴ, accelerates adipogenesis, and produces hypertrophic adipocytes. When challenged with saturated fatty acids, these adipocytes secrete elevated levels of the signature pro-inflammatory adipokines - IL-6, TNF-α, MCP-1, CRP etc., indicating that premature activation of the differentiation program can generate a pro-inflammatory phenotype in adipocytes. Overall, our studies uncovered the regulation of EHMTs and its substantial impact on normal versus pathogenic adipogenesis.

## Results

### Inhibition of EHMT activity expedites adipogenic differentiation in 3T3-L1 pre-adipocytes while Ehmt1/2 over-expression delays it

To investigate how EHMTs are regulated during adipogenesis, we utilized the well-established 3T3-L1 mouse white preadipocyte cell line as the model system. We induced differentiation in 3T3-L1 preadipocytes using the adipogenic cocktail, consisting of isobutylmethylxanthine (IBMX), dexamethasone (Dex) and insulin following the standard protocol (Figure 1A). In the preliminary studies adipogenesis in 3T3-L1 cells were confirmed by the gradual development of intracellular lipid droplets, which appeared as dark red stained granules upon Oil Red O staining (Figure 1B Control panel). In order to study the implications of EHMTs in adipogenic differentiation, we depleted its activity by treating 3T3-L1 preadipocytes with the pan EHMT inhibitor - UNC0642 (27) before adipogenic induction. At Day6 and Day12 time points, accumulation of lipid droplets was found to be markedly increased in control cells compared to DayO (Figure 1B, control panel). However, inhibition of EHMT activity in cells by UNC0642 treatment resulted in increased lipid droplets compared to control cells at Day6 and Day12 (Figure 1B, UNC0642 panel). Intracellular lipid droplets mainly consist of triglycerides (TG), so we measured intracellular TG of differentiating 3T3-L1 cells at different time points DO, D3, D6, D9 and D12. As speculated, TG accumulation increased significantly over time in both control and UNC0642 treated cells. But UNC0642 treated cells showed significantly higher buildup of TG on D6, D9 and D12 by 53%, 46% and 38% respectively compared to untreated control cells (Figure 1C). Hence, this effect was far more pronounced in UNC0642 treated cells vs untreated controls. The augmentation of TG-enriched lipid droplets post-UNC0642 treatment indicate an expedited maturation of adipocytes.

**Figure 1.**
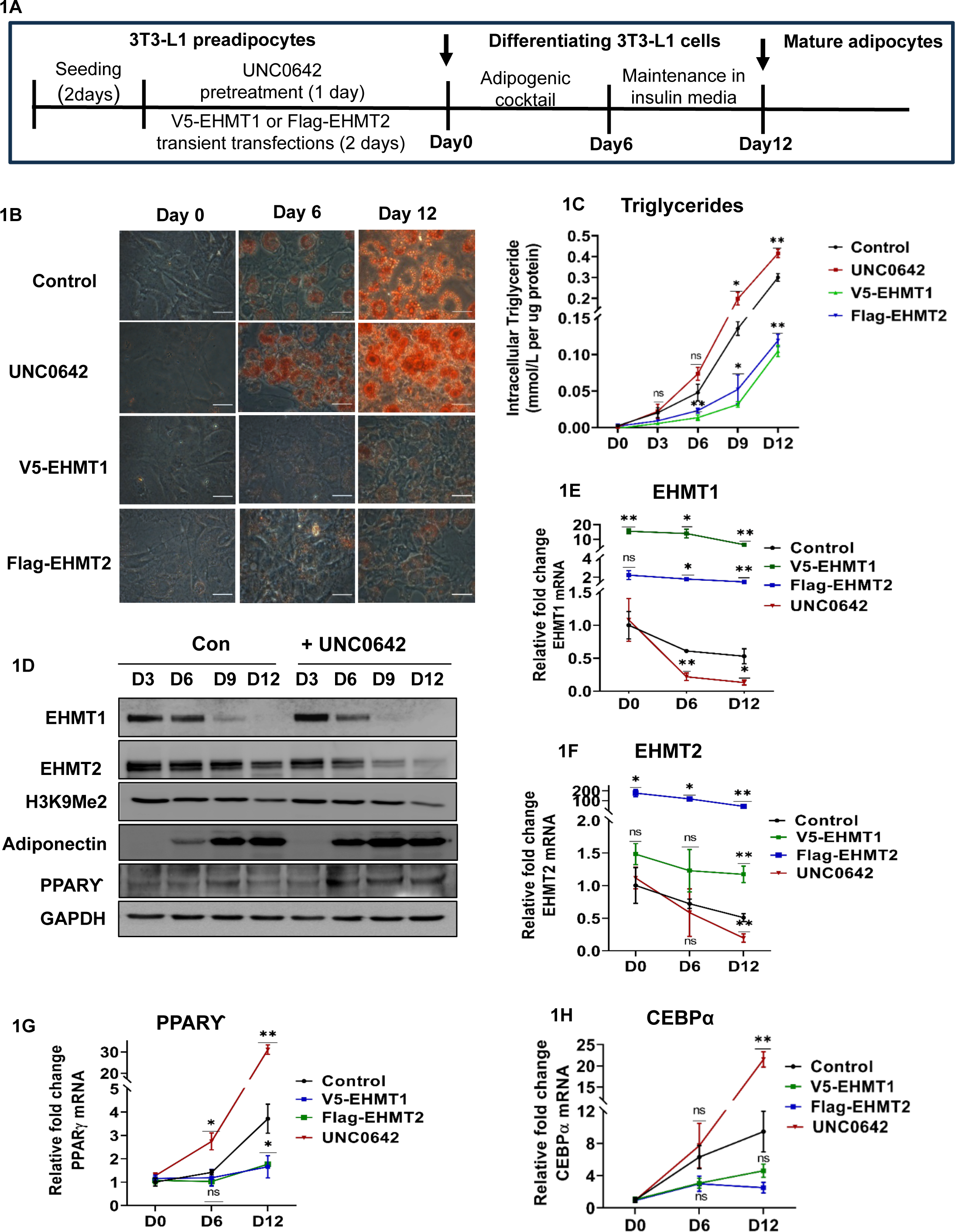
Inhibition of EHMT1/2 activity accelerates adipogenesis, while its over-expression delays adipogenesis. Fig1A: Schema displaying pre-treatment of 3T3-L1 pre-adipocytes with UNC0642 and transient transfections of these cells using *V5-Ehmt1* and *Flag-Ehmt2* constructs, followed by differentiation and collection of cells at different time points for analysis during differentiation. Fig1B: Adipogenic differentiation in Control, UNC0642 treated, V5-EHMT1 and Flag-EHMT2 overexpressed 3T3-L1 preadipocytes as indicated by Oil RedO- stained lipid droplets at Day 0, 6 and 12 after induction of adipogenesis. Fig1C: Intracellular triglyceride accumulation (mmol/L per mg of whole cell lysate protein) in differentiating 3T3-L1 cells with and without UNC treatment, V5-EHMT1 and Flag-EHMT2 over­expression at Day 0, Day 6 and Day 12 post differentiation. Fig1D: Western blot displaying EHMT1, H3K9Me2, Adiponectin, PPARƳ protein expression levels during adipogenesis of 3T3-L1 cells with or without UNC0642 inhibitor pre-treatment. Quantification of protein band intensity analyzed using Image J band densitometry is present in Supplementary FigS1A-S1E. Transcript levels of Fig1E: EHMT1, Fig1F: EHMT2, Fig1G: PPARƳ & Fig1H: CEBPα during the differentiation of 3T3-L1 cells with or without UNC0642 pre-treatment, V5-EHMT1 and Flag-EHMT2 over expression at Day 0, Day 6 and Day 12 post differentiation as analysed by qPCR. Data are representative of three independent experiments. qPCR and Triglyceride quantification data are shown as Mean ± S.D. 2-way ANOVA tests were performed using Prism 8.0 for statistical analyses, p ≤ 0.05 were marked with *, p ≤ 0.01 were marked with ** and considered significant.

Furthermore, to study the individual roles played by EHMT1 and EHMT2, we transfected V5-tagged mouse *Ehmt1* construct *(V5-mEHMT1-pCDNA5/TetO)* and Flag-tagged mouse *Ehmt2* construct *(Flag-mEHMT2-pCDNA5/TetO)* in 3T3-L1 preadipocytes. Post transfection, cells were allowed to differentiate and harvested on DayO, 6 and 12 and analyzed (Figure 1A). *Ehmt1* and *Ehmt2* over-expression impeded adipogenesis as seen by Oil Red O staining (Figure 1B, *V5-Ehmt1* panel and *Flag-Ehmt2* panel). Triglyceride content in *V5-Ehmt1* over-expressed cells was found to be decreased by 71%, 76% and 65% at D6, D9 and D12 time points respectively (Figure 1C) compared to un­transfected controls. While, *Flag-Ehmt2* over-expressed cells led to decrease in TG content by 51%, 61% and 60% at D6, D9 and D12 time points respectively compared to un-transfected controls (Figure 1C).

Interestingly, we noticed that upon adipogenic induction in 3T3-L1 cells, there was a sharp decline in EHMT1 protein levels starting at Day6 in untreated control cells, while EHMT2 protein levels showed minor difference at that time point (Figure 1D) (Figure S1A & S1B). Our data indicates that during normal course of differentiation in control cells, *Ehmt1* transcript (Figure 1E) and protein level (Figure 1D, EHMT1 panel) declines first followed by reduction in *Ehmt2* transcript (Figure 1F) and protein level (Figure 1D, EHMT2 panel). But when methyl transferase activity is inhibited using UNC0642 treatment, both EHMT1 and EHMT2 proteins were reduced at earlier timepoints compared to control while maintaining the trend of sequential downregulation (Figure 1D, UNC0642 panel) (Figure S1A and S1B). This is further supported by decline in the methyl mark - H3K9Me2, specially at D12 (Fig 1D, H3K9Me2 panel) (Figure S1C). In contrast, the master regulator of adipogenesis - *Ppanf* displayed steady and significant increment in its mRNA expression (Figure 1G) by 1.4-fold and 3.7-fold at D6 and D12 respectively as well as increased protein expression (Figure 1D, PPARƴpanel) (Figure S1D) on Day9 and Day12 in untreated control cells. However, UNC0642 treated adipocytes showed 2.7-fold higher *PPARƴ* transcript levels as early as Day6, which increased to a massive 31-fold higher expression at Day12 compared to DayO cells (Figure 1G). Similarly, another principal adipogenic transcription factor CEBPα which is responsible for the mutual expression of *Pparƴ* and itself, showed significant increment in the levels of transcripts by 9.4-fold at Day12 compared to DayO control cells. Upon inhibition of methyltransferase activity, *Cebpα* transcript expression was profoundly altered by a substantial 21-fold increase at Day12 compared to cells at DayO (Figure 1H). On the contrary, over-expression of both *Ehmt1* and *Ehmt2* limited the increase in expression of the master regulators of adipogenesis significantly during differentiation. *Pparf* transcript levels meagerly increased by 1.6-fold in *Ehmt1* over-expressed cells and 1.7-fold in *Ehmt2* over-expressed cells at the end of 12 days compared to DayO cells (Figure 1G). While *Cebpα* transcript levels got elevated by 4.6-fold and 2.5-fold at Day12 in *Ehmt1* and *Ehmt2* over-expressed cells respectively (Figure 1H), indicating that EHMT1 and EHMT2 are individually regulating the expression of the master transcription factors-PPARƴand CEBPα, crucial for adipogenesis. And as a result, adiponectin, which is an adipokine secreted by mature adipocytes was found to be significantly upregulated in UNC0642 treated cells at earlier time points such as Day6 and Day9 compared to untreated controls (Figure 1D, Adiponectin panel) (Figure S1E). This indicates that inhibition of methyltransferase activity in pre­adipocytes accelerates adipogenesis while individual over-expression of *Ehmt1* and *Ehmt2* in restricts the same process.

In order to study the changes in expression of *Ehmt1/2* and *Pparf* at the outset of adipogenic differentiation, we also examined time points as early as 6hrs, 12hrs, Day1, Day3 and Day6. On closer examination of the earlier time points of differentiation, we observed that *Ehmt1* transcript levels declined significantly from D3 onwards (Figure S1F, blue line), followed by a plunge in *Ehmt2* transcript levels at D6 (Figure S1G, blue line). This resulted in significantly increased expression of *Ppanf* at D6 (Figure S1H, blue line), possibly due to depleted methyltransferase activity, contributed by low levels of EHMT1/2, culminating into visible adipogenesis (Figure 1B, D6 Control panel). Therefore, ‘D6’ time point, intriguingly served as a crucial tipping point, at the end of which, both EHMT1 and EHMT2 are sufficiently reduced to relieve the repression on *Ppanf* promoter and initiate latter’s drastic expression. This effect is again expedited by precocious inhibition of methyltransferase activity using UNC0642, shifting the critical time point to 12hrs (Figures S1F, S1G, S1H red lines). Therefore, our data has shown for the very first time that EHMT1 negatively regulates adipogenesis by acting as the primary thrust, which eventually involves EHMT2 in promoting adipogenesis in a temporal manner. Furthermore, loss in EHMT1/2 expedites adipogenesis, while the over-expression of *Ehmt1/2* impairs the same.

### *EHMT1 regulation by PPARƴ-mediated* negative feedback mechanism

Previous studies by *Wang et al.* have shown that during adipogenic differentiation, there is an inverse correlation between the expression of *Pparf* and *Ehmt1/2.* However, we wanted to explore how EHMT1, which is on par with EHMT2 in driving adipogenesis, is regulated during the adipogenic differentiation process. It is known that PPARƴ forms a heterodimer with RXRα and binds to the consensus PPRE (PPAR response element) - (5’-(A/G)GG (T/G)CA (A/G)AG G(T/G)C A-3’) on genes and this heterodimerization appears to be imperative for the DNA binding function of PPARƴ(28)(29). Although PPARƴis critical during terminal differentiation of 3T3-L1 cells and maintenance of mature adipocytes, it is known to be activated within 24 hours of adipogenic induction (30). However, we and other groups have found that its robust expression begins around Day6 and increases thereafter.

To investigate if there exists a feedback regulation between EHMT1 and PPARƴ we pre-treated 3T3-L1 preadipocytes with 1pM RGZ (PPARƴ agonist) and 10μM GW9662 (PPARƴ antagonist) for 24hrs, followed by induction of differentiation using the adipogenic cocktail and studied the relative expression of *Ehmts* and *Pparƴ* (Figure 2A). As expected, RGZ treatment increased adipogenesis while GW9662 reduced the same as seen from Oil Red O staining of mature adipocytes at D12 (Figure 2B). qPCR analysis showed that RGZ treatment increased *Pparƴ* expression by 2-fold and on the other hand GW9662 reduced *Pparƴ* expression by almost 10-fold at the end of 12 days of differentiation (Figure 2C). Interestingly, RGZ treatment led to downregulated expression of both *Ehmt1* (Figure 2D) and *Ehmt2* (Figure 2E) at Day6 and Day12 compared to untreated controls. This clearly indicated that high levels of PPARƴinhibits the expression of the histone methyltransferases *Ehmt1* and *Ehmt2.* This possibly leads to reduced H3K9Me2 repressive marks on adipogenic genes, including *Pparƴ* and drives adipogenesis further on. Conversely, GW9662 which is a selective antagonist of PPARƴ significantly increased *Ehmt1* expression by more than 2-fold (Figure 2D) and *Ehmt2* expression by 1.5-fold (Figure 2E) compared to the untreated controls. Therefore, high amounts of EHMT1/2, as also found in preadipocytes at DayO during normal course of differentiation, maintain Pparƴin repressed state, thereby inhibiting adipogenesis (Figure 2B). Alternatively, upon adipogenic stimulus, EHMT1/2 levels decline, resulting in *Pparƴ* expression. On detailed comparison, we found that upon adipogenic differentiation of 3T3-L1 cells with or without GW9662 treatment, there was a substantial loss in EHMT1 levels as opposed to EHMT2. This led us to primarily focus on regulation of EHMT1.

**Figure 2.**
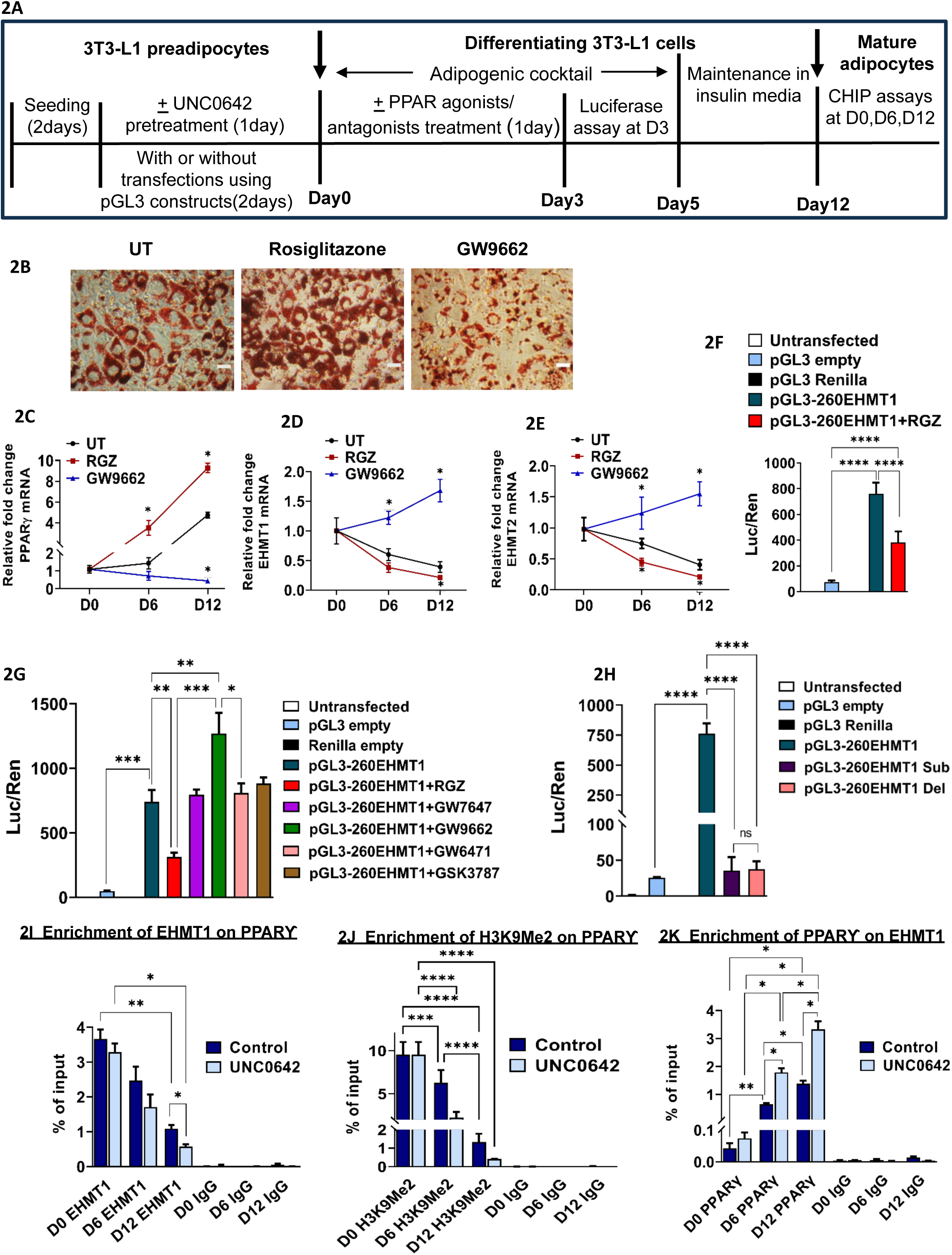
PPARƴbinding to EHMT1 promoter negatively regulates EHMT1 expression. Fig 2A: Schema displaying the timeline for UNC0642 treatment in 3T3-L1 preadipocytes, transfections with different pGL3 & Renilla constructs, PPAR agonists and antagonists treatments, followed by quantification of luciferase activities and CHIP assays. Fig 2B: Oil Red O staining of differentiating 3T3-L1 cells at Day 12, pre-treated with and without Rosiglitazone (PPARƳ agonist) and GW9662 (PPARƳ antagonist). qPCR analysis of Fig 2C-PPARƴ, Fig 2D - EHMT1, Fig 2E - EHMT2 in UT control, RGZ and GW9662 pre-treated 3T3-L1 preadipocytes at various time points during differentiation (DO, D6 and D12). Fig 2F: EHMT1 promoter activity in differentiating 3T3-L1 preadipocytes after transient over­expression of pGL3- 260EHMT1 and pre-treatment with or without PPARƴ agonist - Rosiglitazone (1pM for 24 hrs). Fig 2G: EHMT1 promoter activity of differentiating 3T3-L1 preadipocytes after pre-treatment with various agonists and antagonists of PPAR isoforms after transient over-expression of pGL3-260EHMT1. Fig 2H: 3T3-L1 preadipocytes were transfected with luciferase vector pGL3 carrying 260-EHMT1 promoter fragment with either substitution or deletion mutation and analyzed for EHMT1 promoter activity in terms of luciferase activity. Fig 2I, 2J and 2K: 3T3-L1 preadipocytes were differentiated with or without UNC0642 treatment and cells were collected at DayO, Day6 and Day12. Cell lysates were used to isolate nuclei and chromatin crosslinked to bound proteins. These were chromatin immunoprecipitated (CHIP) with EHMT1, H3K9Me2 and PPARƳ specific antibodies respectively. After reverse crosslinking DNA was isolated and subjected to qPCR and the results were displayed in terms of % enrichment. Data are representative of minimum three independent experiments. Data are shown as Mean ± S.D. 2-way ANOVA with Tukey’s multiple comparison tests and unpaired multiple t-tests were performed wherever needed using Prism 8.0 for statistical analyses, p ≤ 0.05 were marked with *, p ≤ 0.01 were marked with **, p ≤ 0.001 were marked with *** and considered significant.

In order to understand if PPARƴin turn interacts with EHMT1 and assists in completing the possible negative feedback cycle, initiated by EHMT1 itself, we performed in silico analysis using the Cis-bp database. This study revealed that similar to EHMT1 binding to *Pparƴ* promoter sequence, PPARƴ-RXRα heterodimer in turn also binds to the mouse *Ehmt1* promoter region at various sites: - 223 to −240bp, −476 to −489bp, −522 to −528bp, −687 to −699bp, −861 to −867 bp and −944 to −961 bp w.r.t. TSS (Figure S2A). To examine the feedback mechanism in detail, we selected a 260bp fragment from human *Ehmt1* promoter (marked in red letters in Figure S2A), which was shown to contain the consensus PPARƴ-RXRα binding site or PPRE between +146 and +157 bp (marked in blue letters in Figure S2A) of the 600bp long promoter region (Table 2). This 260bp fragment was cloned in pGL3 vector for studying *Ehmt1* promoter activity using luciferase reporter assay. 3T3-L1 cells transfected with the *hEhmt1-P260 pGL3* construct showed increased luciferase activity upon adipogenic differentiation compared to control empty vector transfected cells. But when these cells were treated with Rosiglitazone (PPARƴ agonist), we found a significant reduction in *Ehmt1* promoter activity (Figure 2F). This proved that the increased expression of *Pparƴ* due to agonist treatment resulted in negative regulation of EHMT1 and the existence of a negative feedback pathway. To see if this effect is solely contributed by PPARƴor other isoforms of PPARs like PPARα and PPARβ/δ, we treated the cells harbouring *hEhmt1-P260 pGL3* with various agonists and antagonists of PPAR isoforms and checked for luciferase activity. Interestingly, we found that only PPARƴ agonist RGZ downregulated *Ehmt1* promoter activity, while the antagonist GW9662 upregulated it (Figure 2G). Treatment with other agonist and antagonists of PPARα and PPARβ/δ isoforms didn’t result in any significant difference in promoter activity, ruling out their contribution in this regulation. Furthermore, to test if PPARƴ actually binds to PPRE (PPAR response elements) on *Ehmt1* promoter, we further mutated the binding site by site-directed mutagenesis. We generated a substitution mutation (Figure S2B) and a deletion mutation (Figure S2C) at the PPAR-RXR binding regions on *Ehmt1* promoter using site-directed mutagenesis (SDM) approach. Indeed, 3T3-L1 preadipocytes transfected with mutated constructs (both substitution and deletion mutations) upon adipogenic induction showed negligible *Ehmt1* promoter activity (Figure 2H), indicating that PPARƴ can’t bind to the mutated regions and therefore the binding of PPARƳ to *Ehmt1* promoter was solely responsible for this promoter activity.

**Table 1:**
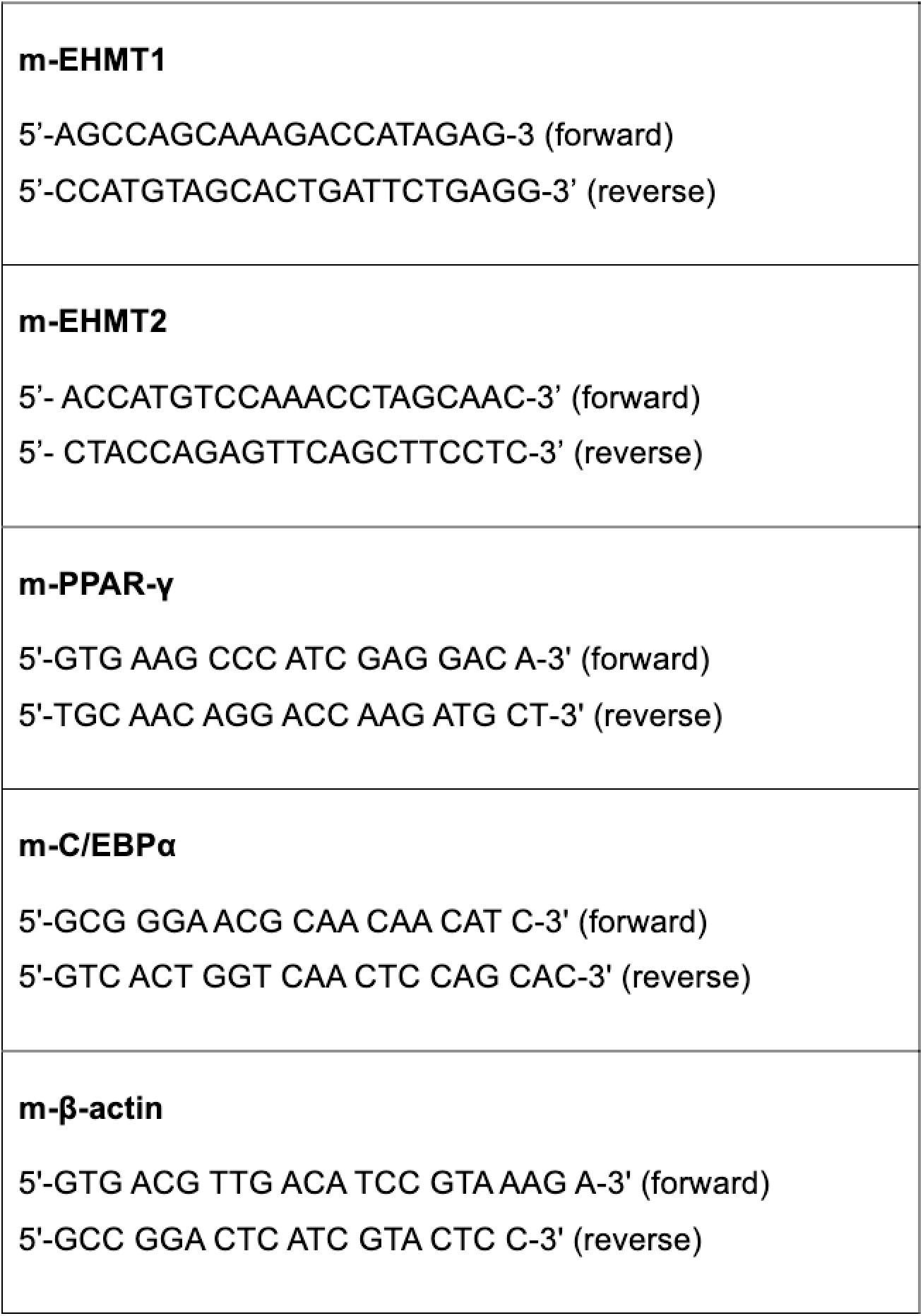
Primers used for qPCR.

**Table 2.**
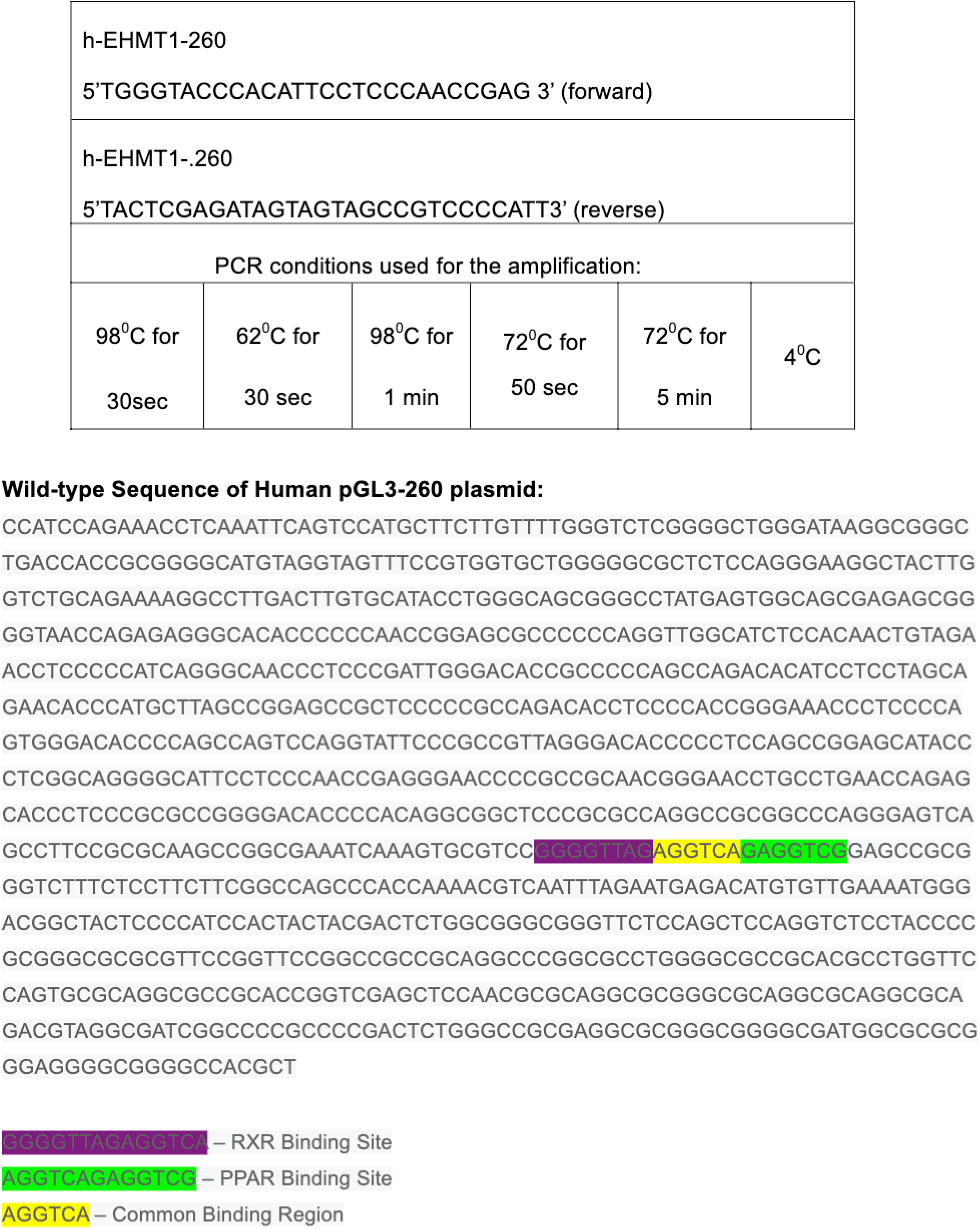
Primers used to amplify human EHMT1 promoter region carrying PPARƴ binding site.

To further reinforce our claim for the existence of a negative regulation, we performed chromatin immunoprecipitation (CHIP) with untreated and UNC0642 treated differentiating 3T3-L1 cells harvested at various time points (DO, D6, D12), using EHMT1 and PPARƴ-specific antibodies for pull down. CHIP-qPCR analysis showed that enrichment of EHMT1 on *Pparƴ* promoter regions is substantially high on DayO (preadipocyte stage) compared to Day 6 and Day12 (Figure 2I). In parallel, H3K9Me2 marks were gradually reduced from Day 0 to Day6 and lowest at Day12 (Figure 2J). Additionally, UNC0642 treatment further reduced EHMT1 enrichment on *Pparƴ* promoter (Figure 2I). Similarly, H3K9Me2 marks were initially enriched on *Pparƴ* promoter and gradually decreased as adipogenesis progressed (Figure 2J). Furthermore, PPARƴ was found to be enriched on *Ehmt1* promoter and significantly increased as the differentiation progressed (Figure 2K). This finally confirmed our claim of the negative feedback regulation.

#### Precocious inhibition of H3K9me2 activity leads to accelerated adipogenesis with inflammatory signatures

Although EHMT1 levels were found to decline strongly as early as Day6 during adipogenic differentiation, there was no substantial loss in its functional activity in terms of H3K9Me2 expression (Figure 1D). Rather a significant reduction in H3K9Me2 levels was found at Day12, which preferably correlated with decline in both EHMT1 and EHMT2 levels at that time point. Therefore, we wanted to investigate if we alter this particular sequence of events by inducing premature loss of global H3K9Me2, what would be the qualitative/quantitative impact on fat or adipokine profile generated by these transformed adipocytes. Indeed, UNC0642 treated cells showed an interestingly unique morphology with lipid droplets coalescing to form large droplets (Figure 1B), which resembled hypertrophic adipocytes found in pathological inflammatory states like obesity and diabetes mellitus. This indicated a possible relationship between early loss of H3K9Me2 and altered fat generation, reflecting inflammation. Clinically, Kleefstra syndrome patients with *Ehmt1* loss of function, gain body weight during childhood and as much as half of them display obesity (19). Moreover, it has been established that obesity is usually associated with low grade chronic inflammation and is responsible for the development of insulin resistance (31). In addition, recent studies have provided strong evidence that exposure to long-chain saturated fatty acids present in high fat diets such as palmitic acid can induce inflammation through TLR4-mediated signalling (32) (33).

To study the link between EHMTs, obesity and inflammation, we therefore treated 3T3-L1 preadipocytes with 200μM of palmitate for 24hrs with or without UNC0642 pre-treatment and with or without over-expressing *V5-Ehmt1* and *Flag-Ehmt2.* At the end of 12 days of differentiation, cells were harvested and extracellular media was collected and stored for further analysis. Cells differentiated in 96-well plate were stained with Oil Red O and imaged using light microscopy. Adipogenesis was found to be highest in UNC0642 treated cells compared to control untreated cells, while *V5-Ehmt1* and *Flag-Ehmt2* over-expressed cells showed the least adipogenesis (Figure 3A, top horizontal UT panel) as earlier. However, upon palmitate challenge the control cells showed mostly unilocular lipid droplets, while *Ehmt1* and *Ehmt2* over-expressed cells showed negligible adipogenesis. Intriguingly, pre-treatment of preadipocytes with UNC0642 along with palmitate led to development of hypertrophic adipocytes, indicative of inflammation (Figure 3A, bottom horizontal PA panel).

**Figure 3.**
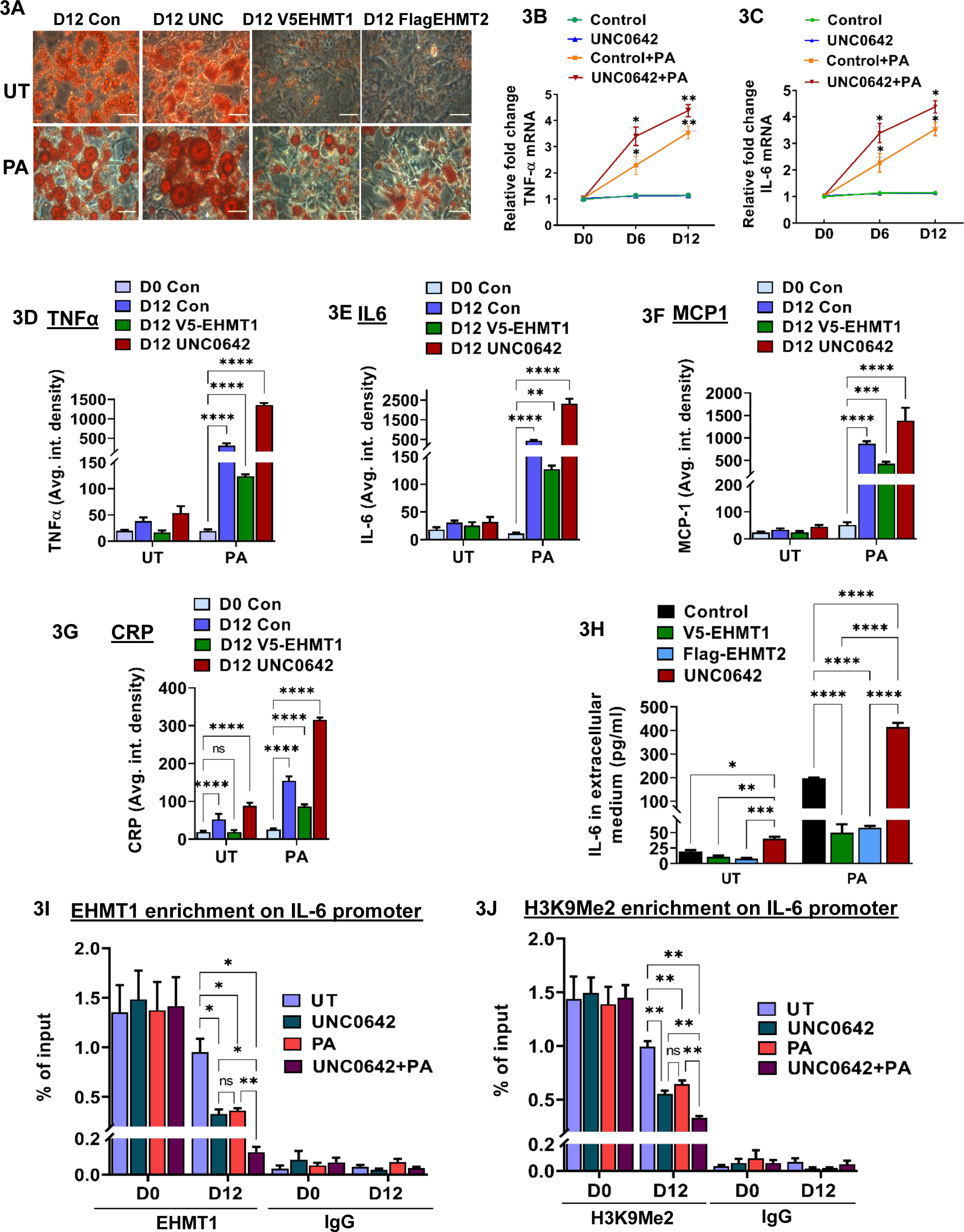
EHMT1 downregulation leads to heightened inflammation upon palmitate challenge. Fig 3A: Oil Red O staining of differentiating 3T3-L1 cells at Day 12, pre-treated with and without 200μM palmitic acid conjugated to BSA and with or without 20nM of UNC0642 for 24 hrs. Fig 3B: TNFα transcript and Fig 3C: IL-6 transcript levels in 3T3-L1 preadipocytes with and without 20nM UNC0642 and 200μM palmitic acid pre-treatment. 3T3-L1 preadipocytes were pre-treated with or without 20nM UNC0642, V5-EHMT1 over-expressed transiently and further treated with or without 200μM palmitic acid for 24 hrs. Cells were differentiated till Day 12 and extracellular media was analyzed for various secreted adipokines using mouse adipokine array. Fig 3D - TNFα, Fig 3E - IL-6, Fig 3F - MCP-1 and Fig 3G - C-reactive protein (CRP). Fig 3H: 3T3-L1 cells were differentiated for 12 days after pre-treatment with or without 20nM of UNC0642 for 24 hrs and V5-EHMT1 transiently over-expressed. Cells were further treated with or without 200μM of palmitate for 24hrs to induce inflammation. Extracellular IL-6 secreted by cells were quantified using indirect sandwich ELISA. Fig 3I: CHIP assay of EHMT1 on IL-6 promoter regions of Control, UNC0642 treated, PA treated and PA+UNC treated 3T3-L1 cells at Day 0 and Day 12 post differentiation. Fig 3J: CHIP assay of H3K9Me2 on IL-6 promoter regions of Control, UNC0642 treated, PA treated and PA+UNC treated 3T3-L1 cells at Day 0 and Day 12 post differentiation. Data are representative of minimum three independent experiments. Data are shown as Mean ± S.D. 2-way ANOVA with Tukey’s multiple comparison tests and unpaired multiple t-tests were performed wherever needed using Prism 8.0 for statistical analyses, p ≤ 0.05 were marked with *, p ≤ 0.01 were marked with **, p ≤ 0.001 were marked with *** and considered significant.

Adipocytes when cultured under *in vitro* conditions also secrete different adipokines in response to various environmental cues. On exposure to saturated free fatty acids, the adipokine profile attains a pro-inflammatory nature. qPCR analysis for the TLR4 downstream inflammatory cytokines - TNFα (Figure 3B) and IL-6 (Figure 3C) revealed that palmitate treated control adipocytes showed gradual increase in expression of both the cytokines at Day6 and Day 12 time points. But when cells are prematurely inhibited for H3K9Me2 activity and then challenged with palmitate, as in UNC0642 treated cells, they showed augmentation in inflammatory characteristics. As a result, UNC0642 and palmitate treated cells showed steep increase in the levels of *Tnfα* and *116* mRNA. Several investigations have shown that the size of the adipocyte is positively correlated to the secretion of the proinflammatory cytokines. So, we checked for secretion of pro-inflammatory cytokines like TNFα, IL-6, MCP-1 and others and quantified them in extracellular media using the mouse adipokine array. We found that when EHMT1/2 activity depleted cells were treated with palmitate, they exhibited hypertrophy and increased amounts of TNFα, IL-6, MCP-1 and C-reactive protein production compared to control cells (Figures 3D, 3E, 3F and 3G respectively). In parallel, we also quantified IL-6 secreted by mature adipocytes at Day12 in the extracellular media using mouse IL-6 ELISA kit. Our data indicate that IL-6 secretion was significantly elevated in cells pre-treated with UNC0642 alone and its levels were further enhanced upon PA treatment (Figure 3H). While, *Ehmt1* and *Ehmt2* over-expressed cells when treated with palmitate showed negligible hypertrophy and attenuated inflammation in terms of IL-6 production (Fig 3D-3G). This confirmed that EHMT1 plays a significant contribution in regulating inflammation as well.

High fat diets (HFDs) consisting of saturated fatty acids are known to signal through TLR2/4 receptors present on adipocyte surfaces and the signal is further transduced through NF-kB pathway, finally activating the downstream inflammatory genes like IL-6, iNOS etc. It has been reported that EHMT1 methylates the promoters of NF-kB target genes, thereby negatively regulating their expression (34). In order to verify this, we investigated if EHMT1 can bind to and deposit methyl marks on the promoter regions of *H6,* which is one of the many pro-inflammatory cytokines produced by adipocytes. To test this, we performed CHIP analysis with untreated control cells and PA treated cells, with or without UNC0642 pre-treatment. Interestingly, we found that *116* promoter is initially enriched with EHMT1 and H3K9Me2 methyl marks in untreated control preadipocytes at DayO. But after 12 days of differentiation, this enrichment is significantly reduced by as much as 2-fold in untreated cells.

While both EHMT1 and H3K9Me2 engagement goes further down on *116* promoter in palmitate or UNC0642 treated cells (Figures 3I & 3J). But when the cells are precociously inhibited of methyltransferase activity, palmitate challenge further reduces the repressive H3K9Me2 marks (Figure 3J) to a minimum, allowing active transcription and translation of *H6* gene, as evidenced by the secreted IL-6 quantified by ELISA in extracellular media (Figure 3H), resulting in a highly inflammatory phenotype. So, if we preprogram the cells very early on by inhibiting histone methyltransferase activity, it leads to adipogenic acceleration accompanied by hypertrophy and secretion of inflammatory cytokines.

#### Downregulation of histone methyltransferase activity in diet-induced obesity mouse model leads to an obese phenotype with heightened inflammation

White adipose tissue (WAT) not only serves as a depot for storage of excess energy but also secretes various bioactive molecules which act in a paracrine fashion to regulate physiological processes, thereby playing a crucial role in energy homeostasis. Excess accumulation of WAT mass results in obesity, which includes both an increase in adipocyte cell size (hypertrophy) and the development of new mature cells from undifferentiated precursors (hyperplasia) (35). Prior studies by *Ohno et al.,* have shown that Adipose-specific *Ehmt1* null mice exhibit obesity and insulin resistance(16). But it was found to be mainly due to reduction in the thermogenic brown fat mass and related abnormalities in metabolism. Our *in vitro* studies indicate that depletion of EHMT1/2 activity using the small molecule inhibitor UNC0642, induces inflammatory signaling in 3T3-L1 pre-adipocytes. And the inflammation is exacerbated in the presence of an adipogenic stimulus like free fatty acids/saturated fatty acids. To test our hypothesis that (a) high fat diet mediated obesity works via EHMT downregulation and (b) UNC0642 treatment could mimic the *in vitro* effects and trigger increased adipogenesis and inflammation *in vivo,* we injected 6-weeks old male C57BL6 mice intraperitoneally with either vehicle (DMSO) or 5mg/kg body weight of UNC0642 prepared in DMSO for 7 consecutive days. Then starting from WeekO, the animals were either put on Chow diet or HFD for a total of 14 weeks. At the end of 14 weeks, all the mice were sacrificed, plasma and adipose tissues were collected for further analysis (Figure 4A).

**Figure 4.**
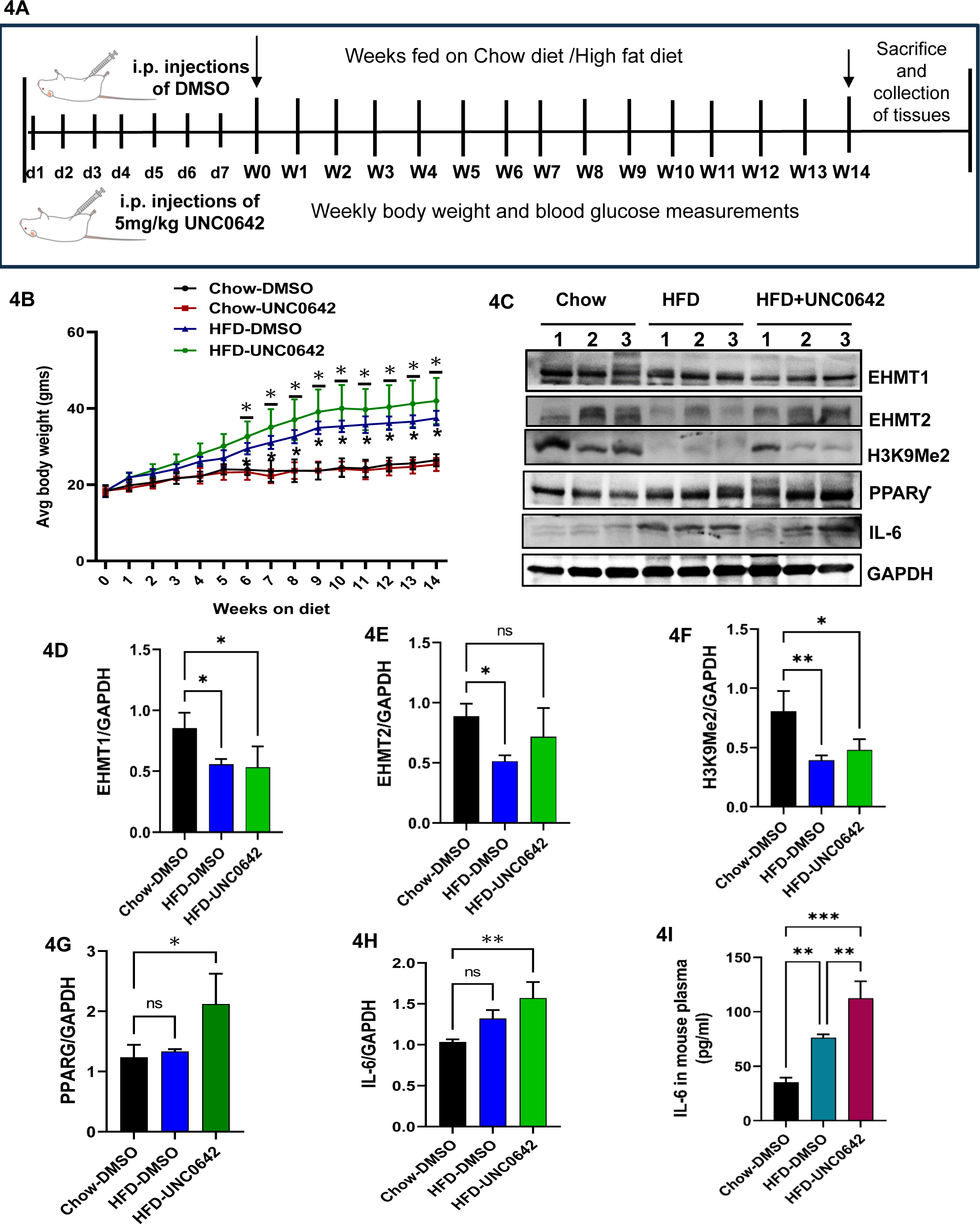
Downregulation of histone methyltransferase activity in diet-induced obesity mouse model leads to an obese phenotype with heightened inflammation. Fig 4A: Schema depicting the timeline for intraperitoneal injection of UNC0642 in 6-weeks old male C57BL6 mice, putting them on respective Chow and High fat diet and further analysis. Fig 4B: Average weekly body weight (in grams) of Chow fed-DMSO injected, Chow fed-UNC0642 injected, HFD fed-DMSO injected and HFD fed-UNC0642 injected mice (n=4 in each group). Fig 4C: Western blot displaying EHMT1, EHMT2, H3K9Me2, PPARƳ and IL-6 protein expression in 50ug total protein from epididymal white adipose tissue (eWAT) of Chow fed-UNC0642 injected, HFD fed-DMSO injected and HFD fed-UNC0642 injected mice (n=3 animal in each group). Fig 4D-4H: Quantification of protein band intensities of Fig4B Western blot analysed using Image J band densitometry. Fig 4I: Plasma IL-6 as quantified by mouse IL-6 ELISA kit in Chow fed-DMSO injected, Chow fed-UNC0642 injected, HFD fed-DMSO injected and HFD fed-UNC0642 injected mice (n=4 in each group). Data are shown as Mean ± S.D. 1-way ANOVA with Tukey’s multiple comparison tests were performed wherever needed using Prism 8.0 for statistical analyses, p ≤ 0.05 were marked with *, p ≤ 0.01 were marked with **, p ≤ 0.001 were marked with *** and considered significant.

**Figure 5.**
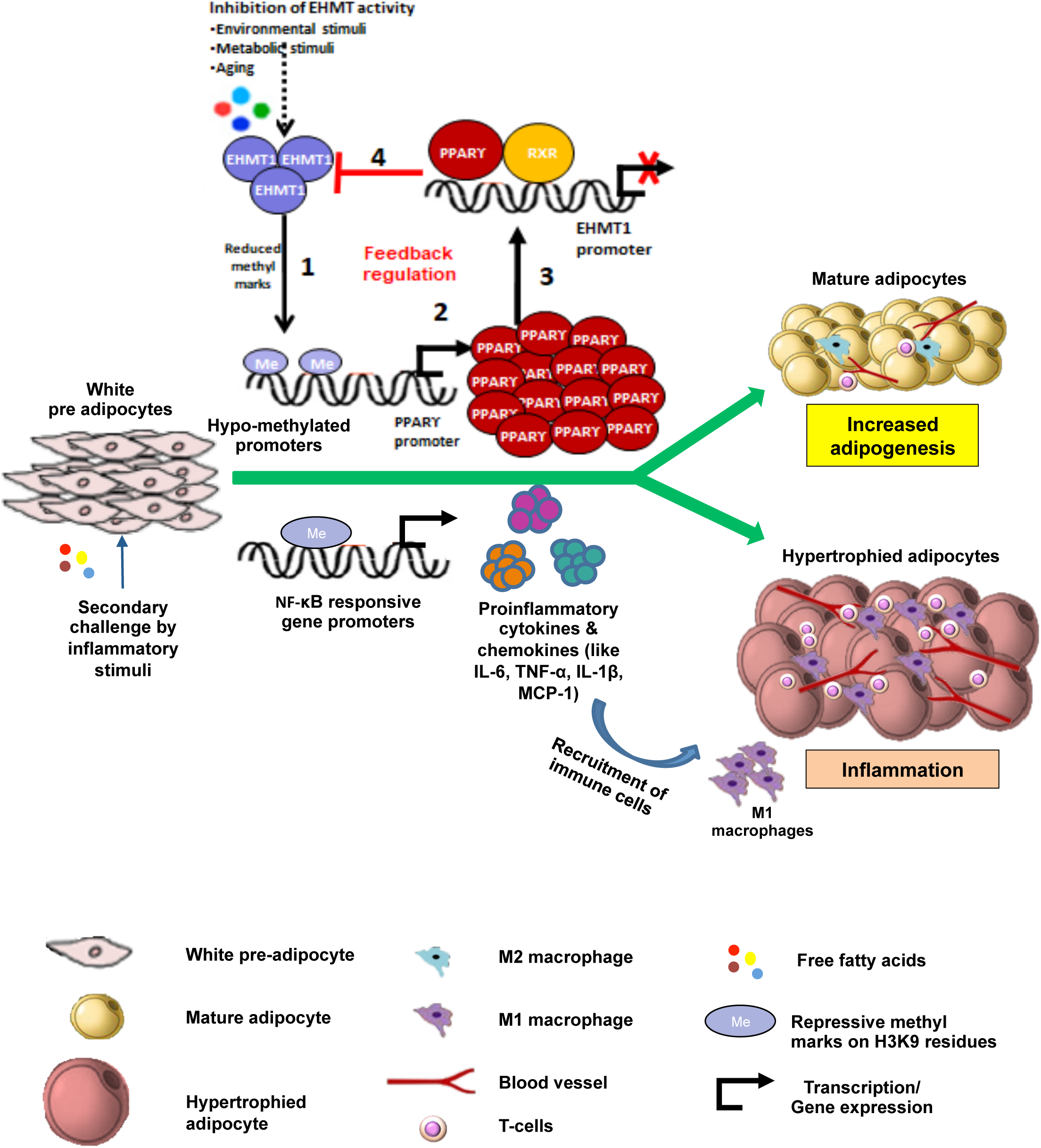
Proposed working model. Initially EHMT1 binds to PPARƴ promoter regions in the preadipocytes, depositing H3K9Me2 repressive marks. But when the cells undergo differentiation, EHMT1 expression declines and PPARƴ expression elevates (Fig 5). Moreover, the increased expression of the demethylase JMJD2B results in reduced enrichment of H3K9Me2/Me3 on PPARƳ and CEBPα promoters, leading to their increased expression and resulting in increased adipogenesis. The large amounts of PPARƴ formed further binds to EHMT1 promoter regions and brings about negative regulation of EHMT1. Thus, EHMT1 expression is sustained at lower level and PPARƴlevels are elevated for adipogenesis to progress.

Unlike the *in vitro* system, wherein UNC0642 pre-treatment, unaided by any other inflammatory stimulus, boosted adipogenesis and hypertrophy in 3T3-L1 cells, we did not observe any increased weight gain in Chow-UNC0642 (UNC0642-injected group) mice compared to Chow-DMSO control mice (Figure 4B). However, high fat diet feeding resulted in accumulation of fat mass and higher body weight, which increased significantly from as early as Week6 till Week14 in both HFD-DMSO (Control) and HFD-UNC0642 (UNC0642-injected group) mice (Figure 4B). Although, it was observed that the HFD-UNC0642 group displayed slightly higher body weight than the HFD-DMSO group, the difference was not significant between the two groups. This could be due to disparity in initial body weights of mice in different groups and also the absorption and bioavailability of UNC0642 in vivo after i.p, administration. It is well established that obesity and insulin resistance go hand in hand. So, we analyzed the mice fasting plasma glucose from Week9 to Week14. Interestingly, blood glucose measurements followed the same pattern as mice body weight, wherein both HFD-DMSO (Control) and HFD-UNC0642 (UNC0642-injected group) mice showed elevated levels of blood glucose compared to Chow diet fed mice with or without UNC0642 treatment (Figure S4A). However, there was no significant difference between the two groups on HFD. Again, we cannot exclude the possibility of variation due to differences in absorption and bioavailability of small molecule (UNC0642).

Upon subjecting the total protein from the eWAT of Chow-DMSO, HFD-DMSO, HFD-UNC0642 mice to Western blot (Figure 4C), we found that HFD fed mice had significantly lower levels of EHMT1 (Figure 4D) and EHMT2 (Figure 4E) compared to Chow diet fed mice. Consequently, the H3K9Me2 repressive marks were significantly reduced in both HFD-DMSO and HFD-UNC0642 mice as compared to Chow-DMSO mice (Figure 4F). In fact, we did not see any additional benefit of UNC0642 in terms of reduction of H3K9Me2 and therefore HFD was enough to alter EHMTs and H3K9Me2. The high fat content of HFD provides an adipogenic microenvironment with the generation of increased levels of PPARƴ (Figure 4G). Moreover, reduced H3K9Me2 marks mediated by lower *Ehmt1/2* expression also contribute to increased expression of PPARƴ and other pro-inflammatory NF-kB responsive genes like *H6* (Figure 4H), *H1 p* and *mcp1.* These mediators further invoke an obesogenic and highly inflammatory milieu resulting in obesity and insulin resistance. Therefore, EHMT1/2 activity can modulate both adipogenicity and inflammation and regulate its own expression via interplay with the master regulator of adipogenesis-PPARƴDuring homeostatic conditions, there exists a balance between the pro- and anti-inflammatory adipokines. However, in the presence of an obesogenic environment, the adipokine profile shifts towards the proinflammatory mediators, promoting inflammation and metabolic disorders related to obesity (36). IL-6 is one such pro-inflammatory cytokine whose concentration is found to be elevated during human obesity and Type-2 diabetes (37). To examine if our diet-induced obese mouse model with UNC0642 treatment (HFD-UNC0642) also exhibit inflammatory characteristics, we subjected plasma from Chow-DMSO, Chow-UNC0642, HFD-DMSO, HFD-UNC0642 mice to ELISA for mouse­specific IL-6 quantification. Intriguingly, we found that HFD feeding resulted in significantly increased IL-6 production. Both HFD-DMSO and HFD-UNC0642 mice plasma showed increase in the amounts of IL-6 in circulation by 2.1-fold and 3.2-fold respectively compared to Chow-DMSO mice plasma, indicating the role played by high-fat diet induced inflammation. However, HFD-UNC0642 mice showed the highest plasma IL-6 concentration, which was significantly higher than the HFD-DMSO control group (Figure 4I), corroborating the pro-inflammatory role played by systemic depletion of EHMT1/2 *in vivo*.

*db/db* obesity mouse model is also known to exhibit increased levels of IL-6 in their circulation (38). Total protein isolated from visceral white adipose tissue (epididymal white adipose tissue) of *db/db* mice when analyzed by Western blot showed almost 20% reduction in EHMT1 protein (Figure S4B and S4C). This further indicated the existence of a negative correlation between EHMT1 and pathogenic adipogenesis as deduced from our earlier *in vitro* experiments.

### Proposed working model

We propose a working model depicting the regulation of EHMT1 during 3T3-L1 preadipocyte differentiation via a dual-axis mode. In the primary axis, an adipogenic stimulus results in decline in *Ehmt1/2* expression, causing hypomethylation of *Pparƴ*promoters, leading to its active expression. This PPARƴ further binds to *Ehmt1* promoters and downregulates *Ehmt1* expression, sustaining adipogenesis while forming a negative feedback regulation. While in the secondary axis, when pre­adipocytes are additionally challenged with an inflammatory stimulus, hypomethylated promoters of inflammatory genes get transformed into active transcriptional states, leading to production of pro-inflammatory milieu. Therefore, EHMT1 regulates itself through interaction with PPARƴ and drives adipogenesis as well as inflammation during adipogenesis.

## Discussion

Adipogenic differentiation proceeds through a sequential cascade of events involving several transcription factors and chromatin modifying co-regulators acting via various signaling pathways (20). Although, it is known that both EHMT1 and EHMT2 are essential in regulating adipogenesis (16)(25), very little is known about their individual contributions or concerted action and precise temporal expression during adipogenesis. Moreover, it’s not clearly understood how these epigenetic regulators are regulated themselves. Here we report that regulation of adipogenesis by EHMTs is indeed achieved by a two-step process, involving the sequential downregulation of *Ehmt1* followed by that of *Ehmt2* and a mutual interaction with PPARƴ, resulting in sustained adipogenesis. Our studies deciphered that while the downregulation of *Ehmts* occur during the initiation phase of a typical adipogenic differentiation program, the methyltransferase activity is diminished during the subsequent maturation phase, when levels of both EHMT1 and EHMT2 are sufficiently reduced. This results in facilitating *Pparƴs* steady expression leading to successful adipogenesis. However, if we skew the methyltransferase activity either by inhibiting enzymatic activity or over-expression of *Ehmt1/2,* it leads to accelerated or blunted adipogenesis respectively. Altogether, these results clearly indicate the proficiency of EHMT1 in regulating adipogenesis; in which EHMT1 plays a crucial role during the initial stages, while EHMT2 plays a concerted role along with EHMT1 at the later stages of adipogenesis. We presume this dictating role of EHMT1 is because of its inherent capacity to function both as an epigenetic reader and a writer. In addition, it is also possible that this governing role of EHMT1 in adipogenic differentiation could be due to its ability to stabilize the EHMT1-EHMT2 complex in order to function as a holoenzyme.

The dynamic state of histone lysine methylation is balanced by activities of both histone methyltransferases and histone demethylases (21)(39)(40). The H3K9Me2/Me3 demethylase - JMJD2B is known to promote adipogenesis by reducing the enrichment of H3K9Me2/Me3 on the promoters of both *Ppaiy* and *Cebpα* (41)(21). In line with this study, we speculate that during the initial phase of adipogenic induction, *jmjd2b* expression gradually gets upregulated in order to augment the removal of repressive H3K9Me2/Me3 marks from *Ppaiy* and *Cebpα* promoter regions, stimulating the expression of the master regulators of adipogenesis. PPARƴ then acts on *Ehmt1* to initiate the sustainable and robust adipogenic signalling. While our study demonstrates that EHMT1 causes the primary thrust in the initiation of adipogenesis, we cannot completely rule out the possible contributions of EHMT2 in this process. The question as to what marks the PPARƴ activation during very early stages of adipogenesis needs further investigation.

Ever so, PPARƴ can be activated by agonists like insulin-sensitizing thiazolidinediones drugs (eg. rosiglitazone), resulting in robust adipogenic differentiation (28). Interestingly, we found that RGZ downregulated expression of both *Ehmt1 and Ehmt2* and upregulated *Pparƴ*expression, leading to enhanced adipogenesis. Conversely, GW9662 treatment upregulated both *Ehmt1 and Ehmt2* expression and downregulated *PparƳ*expression, leading to inhibition of adipogenesis. These findings indicated for the very first time that there exists a feedback regulation between EHMT1/2 and PPARƴ through which EHMT1 levels are regulated during adipogenesis. Precisely, our studies reveal that during the pre-adipocyte stage, EHMT1 binds to *PparƳ*promoter regions, deposits H3K9Me2 marks, maintaining PPARƴin repressed state. However, in the presence of an adipogenic stimulus, *Ehmt1* expression declines, depositing lesser H3K9Me2 marks on the promoter of *PparƳ,* leading to active expression of the latter. The constantly increasing levels of PPARƴ further binds to *Ehmt1* promoter regions and downregulates the latter’s expression. Thus, the cells attain a steady state, wherein *Ehmt1* expression is maintained at extremely low levels, while *PparƳ*expression is highly elevated for adipogenesis to continually progress further.

Excess calorific intake, ageing, genetic and epigenetic changes along with environmental factors trigger obesity in White Adipose Tissue (WAT), characterized by adipocyte dysfunction, hypertrophy, ectopic lipid accumulation, and insulin resistance (42). Free fatty acids released from excess lipid-laden hypertrophied adipocytes can induce TLR4 signalling via NF-kB activation (43)(37), thereby inducing homeostatic inflammation during obesity (31)(44). Prior studies have shown that EHMT1 inhibits the expression of inflammatory genes by depositing H3K9Me2 marks on the NF-kB responsive gene promoters (34). Our data indicate that pre-adipocytes additionally challenged with an inflammatory stimulus like saturated fatty acids, results in hypertrophic adipocytes with inflammatory fat. And this phenotype gets magnified when H3K9Me2 activity is precociously inhibited in the presence of saturated fatty acids. The hypomethylated promoters of NF-kB responsive inflammatory genes switch into active expression states. This leads to upregulation of inflammatory genes and expression of proinflammatory mediators like IL-6, IL-1 p, MCP-1 and MCSF-1. We found that while loss of EHMT1/2 activity exacerbated the inflammatory phenotype, over-expression of *Ehmt1/2* mitigated inflammation. *In vivo* studies using the diet-induced obesity mouse model also confirmed this finding. High fat diet triggered increase in body weight in mice but UNC0642 treated mice on high fat diet displayed a propensity towards higher weight gain and systemic inflammation compared to control mice. As adipogenesis goes hand in hand with angiogenesis, increased production of the adipokine - VEGF stimulates the propagation of new blood vessels. This neo-angiogenesis is believed to further aid in transporting immune cells like monocytes and T-cells to the adipogenic milieu, generating elevated levels of inflammatory mediators like IL-6, transforming the healthy adipose tissue into a hypertrophic, metabolically unstable and highly inflamed adipose tissue. Thus EHMT1/2 can modulate the inflammatory signature of adipocytes. However, it is not clear yet, if there exists similar negative feedback regulation between EHMT1 and PPARƴ during pathogenic adipogenesis. Taken together, our studies have systematically uncovered the regulation of chromatin modifiers - specifically EHMTs - and their activities during adipogenic differentiation, and established that skewing the fine balance of this regulatory mechanism tips the homeostatic process towards a pathogenic inflammatory state.

## Materials and methods

### Cell culture

3T3-L1 preadipocytes were grown to confluence in growth medium consisting of high-glucose DMEM supplemented with 10% fetal bovine serum and 100 U/mL penicillin and 100 μg/mL streptomycin. Two days post-confluence, differentiation medium consisting of growth medium supplemented with 1pM dexamethasone, 10 μg/mL human insulin, and 0.5 mM isobutyl-1-methyl xanthine (IBMX) was added. Cells were grown in differentiation medium for 6 days, followed by 6 days in growth medium with insulin. The differentiation medium was replaced on alternate days.

### Oil red O staining

3T3-L1 adipocytes were washed twice with phosphate-buffered saline (PBS), fixed for 1 h with 4% formaldehyde in PBS, and subsequently dehydrated with 60% isopropanol (Amresco, OH) for 5 min. After removing the isopropanol completely and drying the wells, Oil Red O stain diluted with DDW was added to the plate, which was incubated for 10-15 mins. Subsequently, Oil Red O was removed and excess dye was washed off 4 times with distilled water until the background was clear. Images were obtained using an Evos Original microscope (Advanced Microscopy Group).

### RNA isolation, qPCR, and gene expression profiling

RNA was isolated using Trizol method, followed by reverse transcription of 1μg of RNA with cDNA Reverse Transcription kit (Invitrogen) following manufacturer’s instructions. qPCR was performed using primers as described (Tablel), SYBR Green PCR Mastermix (Invitrogen), and the CFX 384 instrument (Biorad). Analysis was performed using the delta Ct method and normalization of all genes of interest to the housekeeping control p-actin.

### Immunoblotting

Cell protein was extracted on ice in cold whole-cell RIPA buffer supplemented with protease inhibitor cocktail (Roche). SDS-PAGE was performed using 10% Tris-glycine gels, followed by transfer to PVDF membranes (In-vitrogen). The primary antibodies used for immunoblotting were as follows: anti-EHMT1, anti-EHMT2, anti-PPARƳ, anti-H3K9Me2, anti-adiponectin. After incubation with HRP-conjugated secondary antibodies, blots were developed using the enhanced chemiluminescent substrate kit (Invitrogen).

### Plasmids, transfections, and luciferase reporter assays

Human EHMT1 promoter sequence was retrieved using Eukaryotic Promoter Database (EPD). The catalog of preferred binding sequences was found using CIS-BP database, where the PPARƴ and RXRα binding sites were found to be present at ((+146) to (+157)) position relative to the Transcription Start Site (TSS). A ∼260-bp enriched region was amplified from human genomic DNA using PfuTurbo Hotstart Polymerase (Stratagene) using primers listed in Table 2 and cloned into the pCR-Blunt ll-TOPO vector (Invitrogen). Following restriction enzyme digest, the regions were inserted into the multiple cloning site of the pGL3-Promoter vector (Promega). 3T3-L1 cells were transfected overnight with 1μg of pGL3-Promoter construct, 50 ng of pRL-CMV renilla vector or 1μg of pGL3 empty in 24-well plates using Lipofectamine LTX plus (Invitrogen) according to manufacturer’s protocol. Luciferase activity was measured using Dual Reporter Assay (Promega), normalizing firefly luciferase to renilla activity.

### Site directed mutagenesis

Following parameters were used for PCR amplification of H260-PGL3 plasmid: 98°C for 30 Sec, 18 cycles (98°C for 10 s, 70°C for 30 s, 72°C for 3 min), and 72°C for 7 min in a final reaction volume of 25pl. The reaction product was digested with 10 U of methylation­sensitive enzyme Dpnl at 37°C for 1 h. (R0176, New England Biolabs). E. coli DH5-a competent cells were transformed with the amplified products. Finally, the plasmids were purified using the Promega Wizard Plus SV Minipreps DNA Purification Systems. The sequences of the primers used for mutation is provided in Table 3. The mutated plasmids were subjected to Sanger sequencing and sequence of the mutated region in the H260-PGL3 plasmid is provided in Table 4.

**Table 3.**
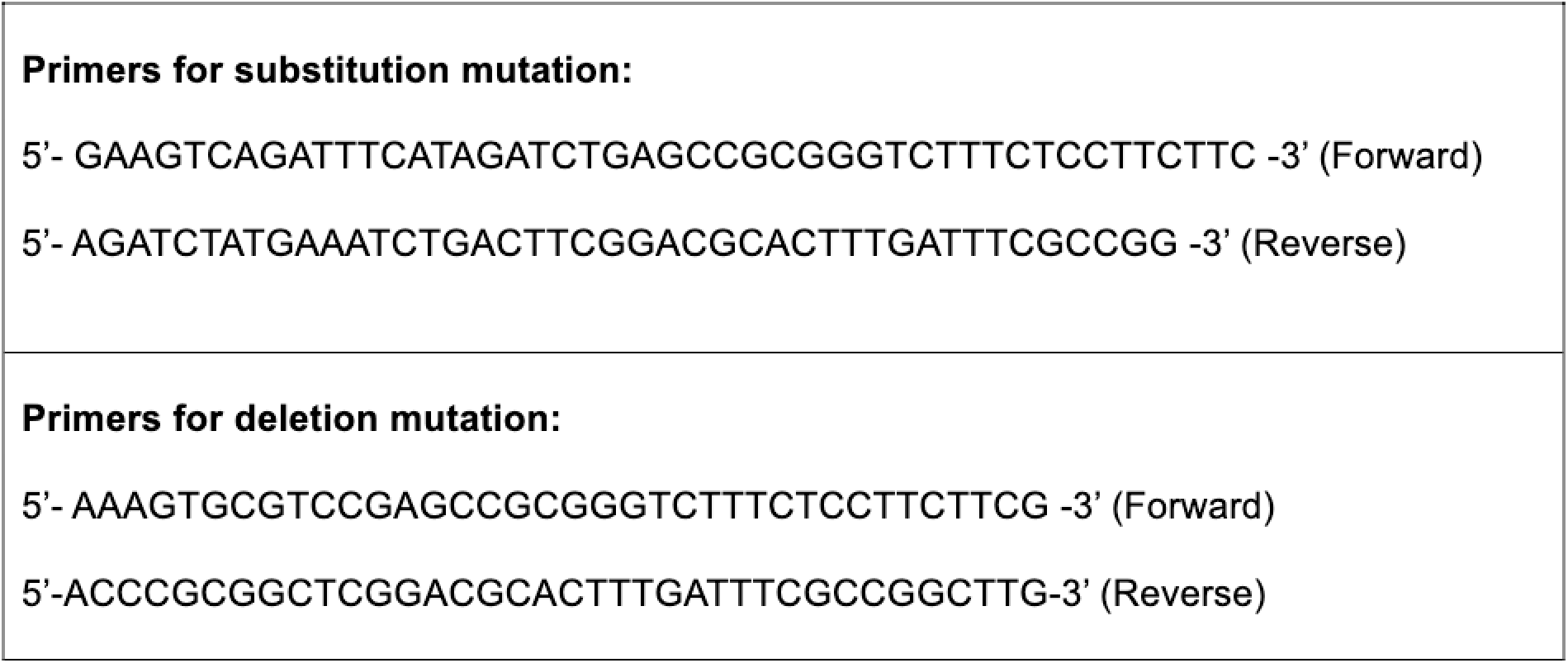
Primers used for site-directed mutagenesis.

**Table 4.**
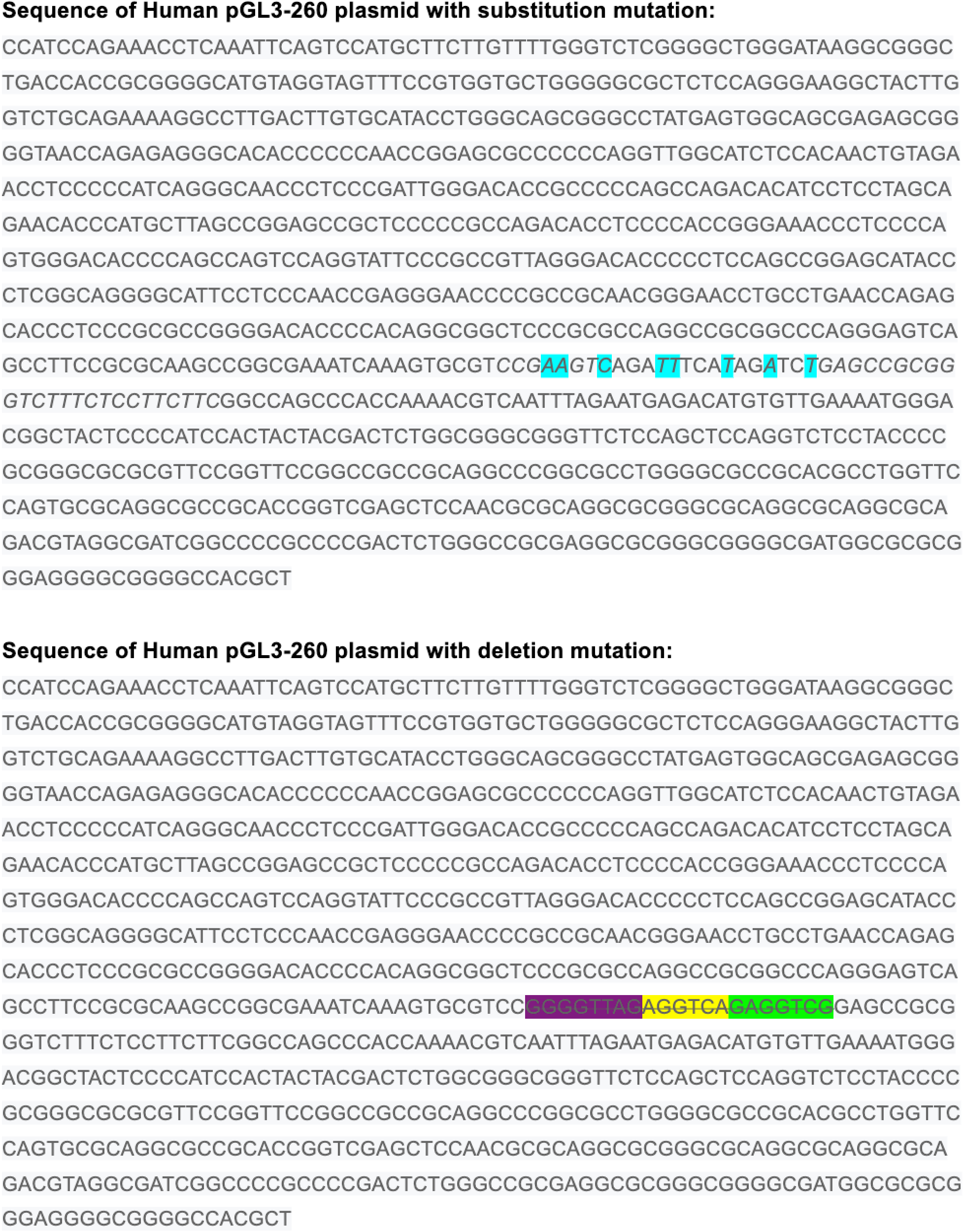
Sequence of the mutated region in the H260-PGL3 plasmid.

### ChlP

Cells were cross-linked in 1% formaldehyde for 15 mins, followed by quenching with 1/20 volume of 2.5 M glycine solution, and two washes with 1* PBS. Nuclear extracts were prepared by dounce homogenizing in nuclear lysis buffer (20mM HEPES, 0.25Msucrose, 3 mM MgCI2, 0.2% NP-40, 3mM p-mercaptoethanol, 0.4mMPMSF, Complete protease inhibitor tablets from Roche). Chromatin fragmentation was performed by MNase treatment in ChlP SDS lysis buffer (50mMHEPES, 1% SDS, 10 mM EDTA). Proteins were immunoprecipitated in ChIP dilution buffer (50mMHepes/NaOH at pH 7.5, 155 mM NaCI, 1.1% Triton X-100, 0.11% Na-deoxycholate, 1mM PMSF, Complete protease inhibitor tablet), using nonspecific rabbit IgG control and mouse IgG control (Sigma). Cross-linking was reversed overnight at 65°C, and DNA isolated using phenol/chloroform/isoamyl alcohol. For ChlP-qPCR, enrichment was measured using SYBR Green PCR Mastermix and the Biorad CFX38 instrument (Biorad). Analysis was performed by the % input method. Primer sequences used for qPCR analysis can be found in Table 5.

**Table 5.**
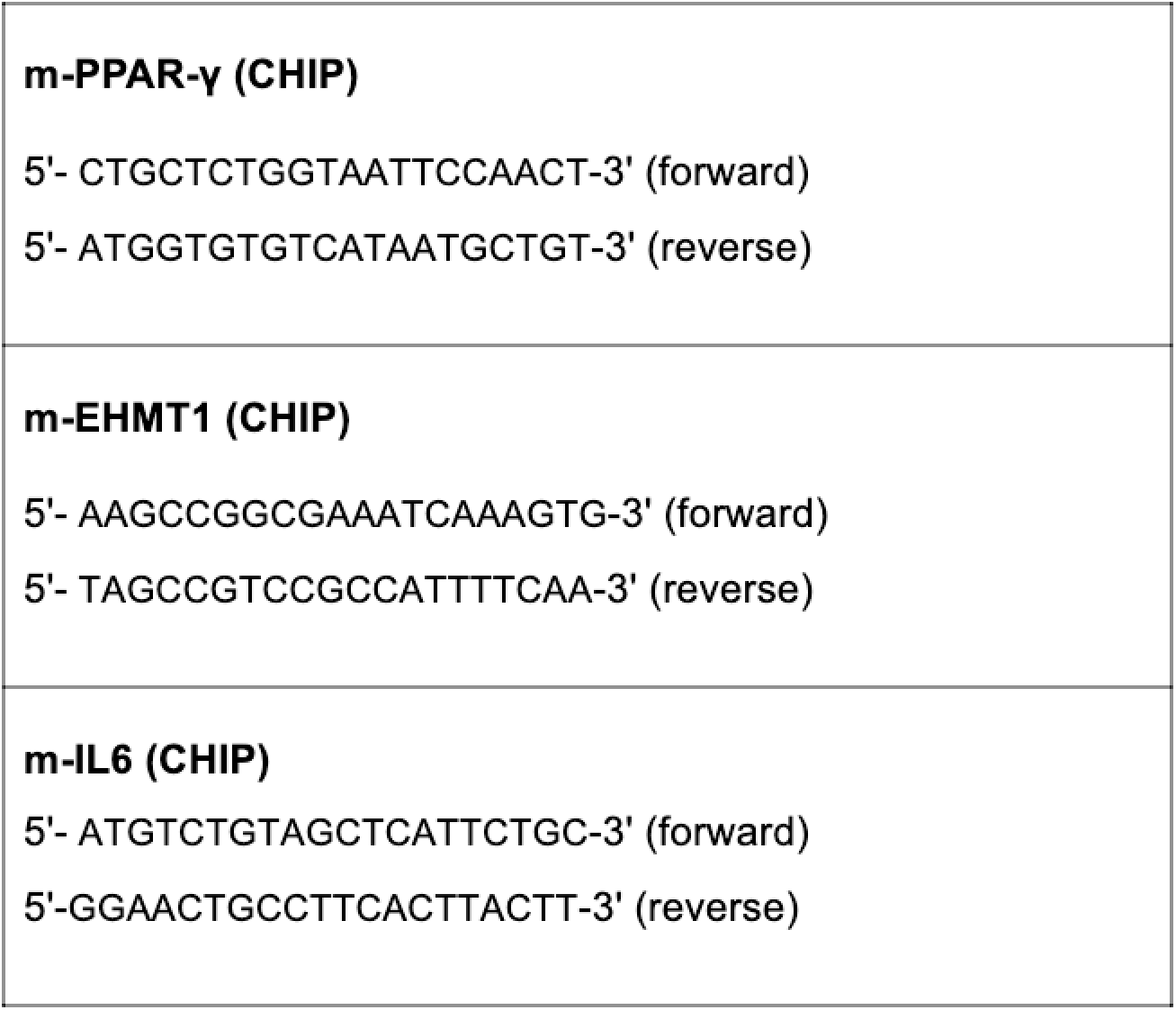
CHIP qPCR mouse primer sequences.

### Palmitate treatment

Free fatty acid in the form of palmitic acid (sodium-palmitate; Sigma-Aldrich) were conjugated with FFA-free bovine serum albumin (BSA) for administration to cells. Briefly, FFAs were dissolved in ethanol and diluted in DMEM containing 1% FBS and 2% (wt/vol) BSA for 1hr at 55°C. BSA-conjugated FFA-containing media were used to challenge cells.

### Mouse Adipokine array

Extracellular conditioned media from differentiated 3T3-L1 cells, over­expressed with or without V5-mEHMT1 and pretreated with or without palmitate and UNC0642 were quantified for 38 different adipokines using Proteome Profiler Mouse Adipokine Array Kit ARY013 from R&D Systems as per the manufacturer’s protocol. Dot blot analysis was carried out using Image J software.

### ELISA

Extracellular conditioned media from differentiated 3T3-L1 cells pretreated with or without palmitate and UNC0642 were quantified using mouse IL-6 Quantikine ELISA kit M6000B from R&D Systems as per the manufacturer’s protocol. Mouse plasma was also assayed for IL-6 levels using the same method.

### Animal handling and procedures

6 weeks old male C57BL6 mice were used for all the animal work. Animals were maintained at SPF2 Animal facility in NCBS/inStem Animal Care and Resource Center (ACRC) with fixed light and dark cycles, controlled humidity and temperature and fed with Chow or High fat diet at *ad libitum.* For the UNC0642 treatment studies, 6-weeks old males were injected intraperitoneally with either DMSO or 5mg/kg body weight of UNC0642 prepared in DMSO for 7 consecutive days. Then starting from Week 0, their body weights were measured. At the end of 14 weeks, all the mice were sacrificed, adipose tissues and plasma samples were collected and snap frozen for further analysis and stored at −80degC. Epididymal white adipose tissue (eWAT) were homogenized in RIPA buffer containing protease inhibitors and equal amounts of proteins were subjected to Western blot analysis. Plasma samples were analyzed for II-6 levels using mouse IL-6 ELISA. All animal procedures were operated and approved according to the Instem or NCBS IAEC under IAEC project# NCBS-IAE-2018/02(N).

## Author contributions

MC designed and performed experiments on adipogenesis, UNC0642 and palmitate treatment and over-expression studies, Oil Red O staining, qPCRs, Western blots, luciferase assays, ELISA, Adipokine arrays, Chromatin immunoprecipitation studies, performed statistical analyses, assembled the figures and wrote the manuscript. JA is responsible for cloning the hEHMTI promoter-pGL3 constructs and VKK performed the site-directed mutagenesis in these hEHMTI-pGL3 clones. NS intraperitoneally injected the C57BL6 animals with UNC0642, put them on High fat diet, measured weekly body weights and blood glucose of mice and collected tissues at the end of 14 weeks. SR conceptualized the project, designed the experiments, helped with the manuscript writing and guided the entire project.

## Acknowledgements

This work was supported by core funding provided to S.R. from inStem. M.C. acknowledges Department of Science and Technology, Government of India for receiving financial support vide reference number SR/WOS-A/LS-221/2017 under the Women Scientist Scheme-A postdoctoral fellowship to carry out this work. VK is supported by UGC-CSIR Junior Research Fellowship (JRF). The authors acknowledge the facilities, and the scientific and technical assistance of members from Dr. Colin Jamora lab at the Institute of Stem Cell Science and Regenerative Medicine (inStem), Bangalore. Animal work in the NCBS/inStem Animal Care and Resource Center was partially supported by the National Mouse Research Resource (NaMoR) grant # BT/PR5981/MED/31/181/2012; 2013-2016 & 102/IFD/SAN/5003/2017-2018 from the Indian Department of Biotechnology.

## Conflict of interest

The authors have declared that no conflict of interest exists.

## Supplementary Figure Legends

**Supplementary Figure 1:**
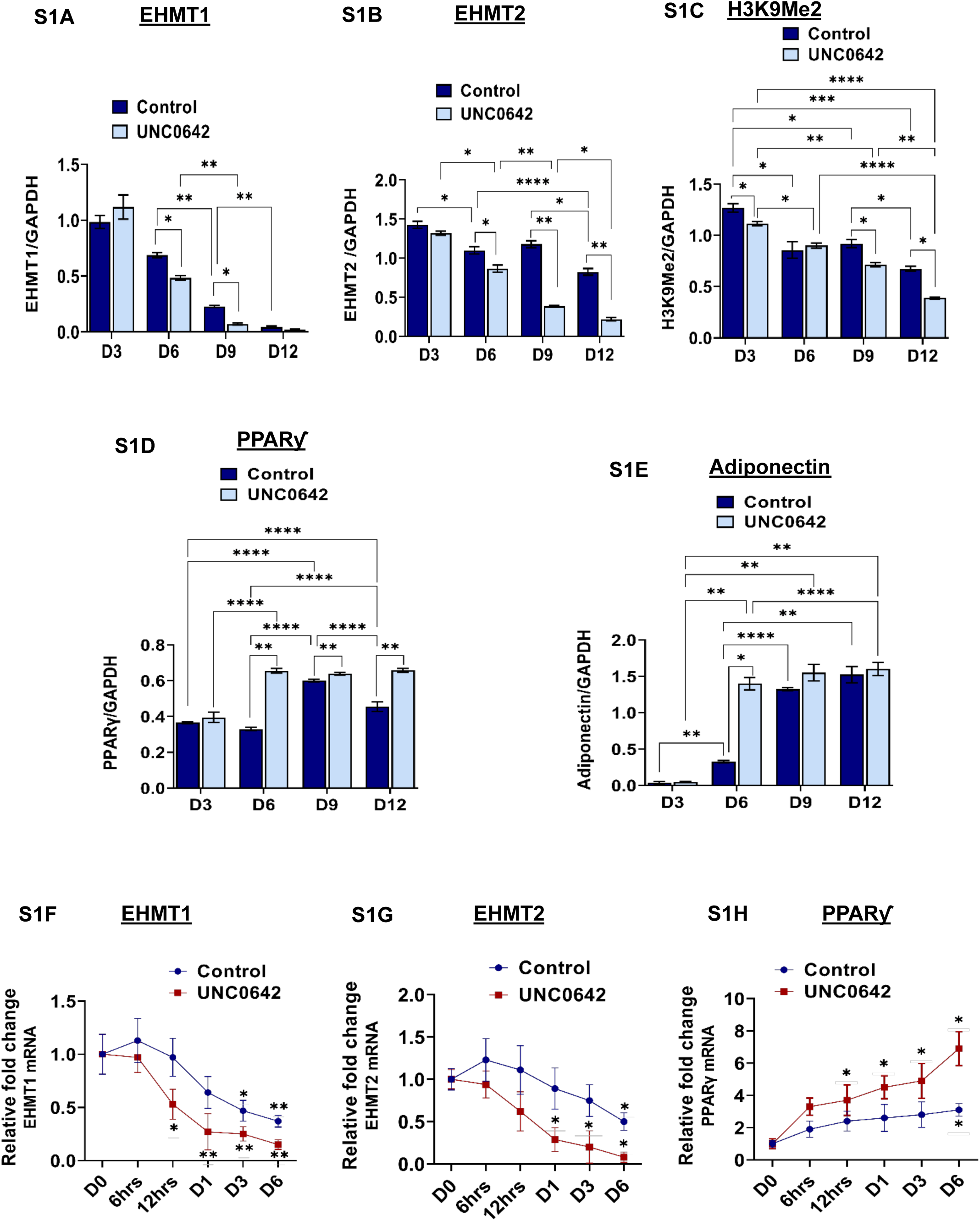
Supp Fig S1: Band densitometry for Western blot performed using whole cell lysates from Control and UNC0642 treated 3T3-L1 cells at different days of differentiation (‘D’ indicates days post differentiation). Image J software was used for analysing band densities and were normalized with respective GAPDH band densities. S1A- EHMT1, S1B - EHMT2, S1C - H3K9Me2, S1D - PPARƴand S1E - Adiponectin. Supp Fig S1F - transcript levels of EHMT1, Fig S1G - transcript levels of EHMT2 and Fig S1H - transcript levels of PPARƴin differentiating 3T3-L1 preadipocytes pre-treated with or without UNC0642 were collected at early time points - Ohr, 6hrs, 12 hrs, D1, D3 and D12 post differentiation. Data are representative of minimum three independent experiments. Data are shown as Mean ± S.D. 2-way ANOVA with Tukey’s multiple comparison tests and unpaired multiple t-tests were performed wherever needed using Prism 8.0 for statistical analyses, p ≤ 0.05 were marked with *, p ≤ 0.01 were marked with **, p ≤ 0.001 were marked with *** and considered significant.

**Supplementary Figure 2:**
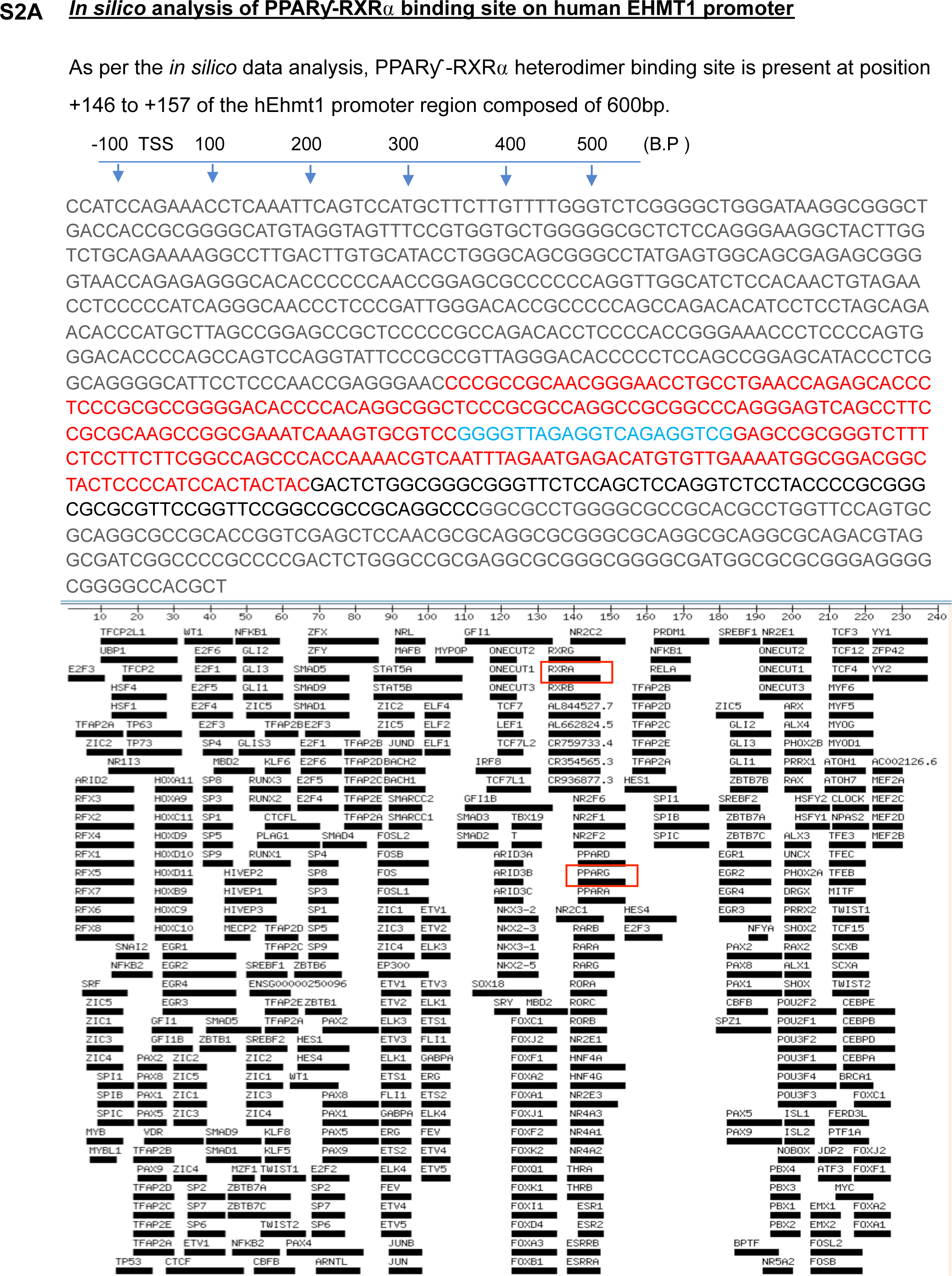
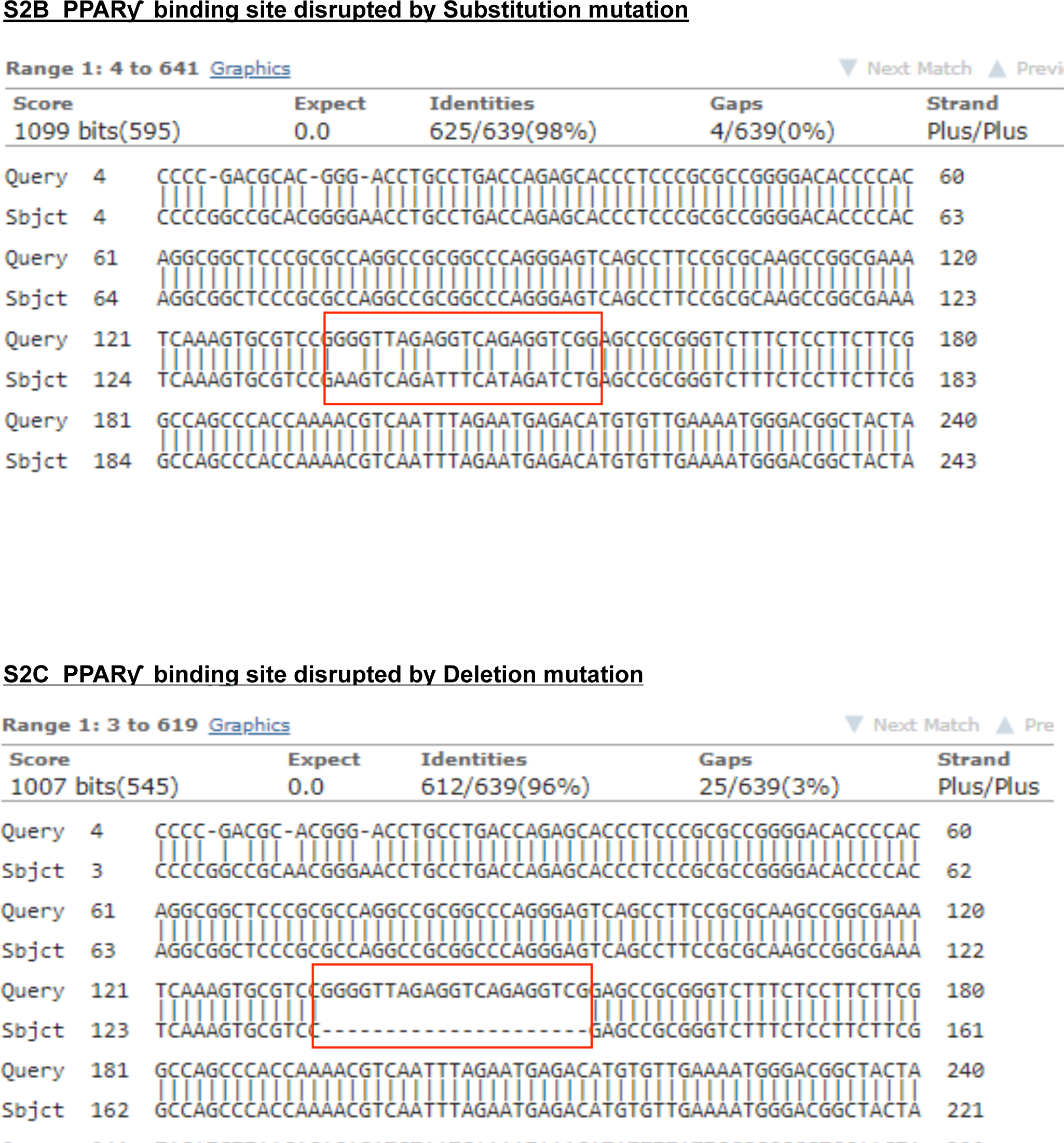
Supp Fig S2A: *In silica* analysis of binding sites for PPARƳ-RXRα on human *Ehmt1* promoter using cis-bp database. Supp Fig S2B: PPARƴ binding site on *hEhmt1* promoter disrupted by substitution mutation. Supp Fig S2C: PPARƴ binding site on hEHMT 1 promoter disrupted by deletion mutation. Data are representative of minimum three independent experiments. Data are shown as Mean ± S.D. 2-way ANOVA with Tukey’s multiple comparison tests and unpaired multiple t-tests were performed wherever needed using Prism 8.0 for statistical analyses, p ≤ 0.05 were marked with *, p ≤ 0.01 were marked with **, p ≤ 0.001 were marked with *** and considered significant.

**Supplementary Figure 3:**
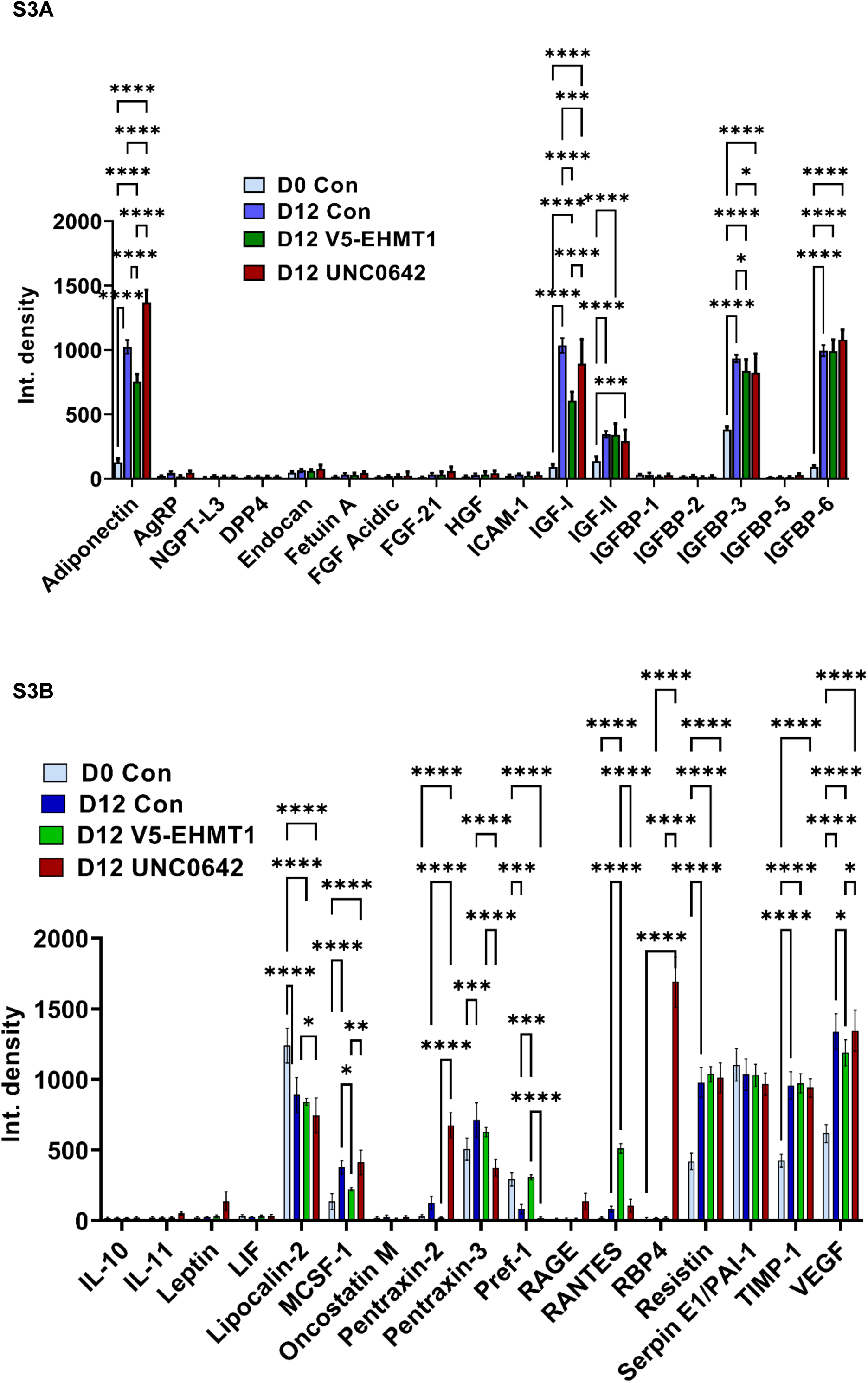
Supp Fig S3A and S3B: Amount of adipokines secreted by Untreated control, PA+Control, PA+UNC0642 treated, PA+V5-EHMT1 overexpressed 3T3-L1 preadipocytes at Day 0 and at Day 12 post differentiation. Data are representative of minimum three independent experiments. Data are shown as Mean ± S.D. 2-way ANOVA with Tukey’s multiple comparison tests and unpaired multiple t-tests were performed wherever needed using Prism 8.0 for statistical analyses, p ≤ 0.05 were marked with *, p ≤ 0.01 were marked with **, p ≤ 0.001 were marked with *** and considered significant.

**Supplementary Figure 4:**
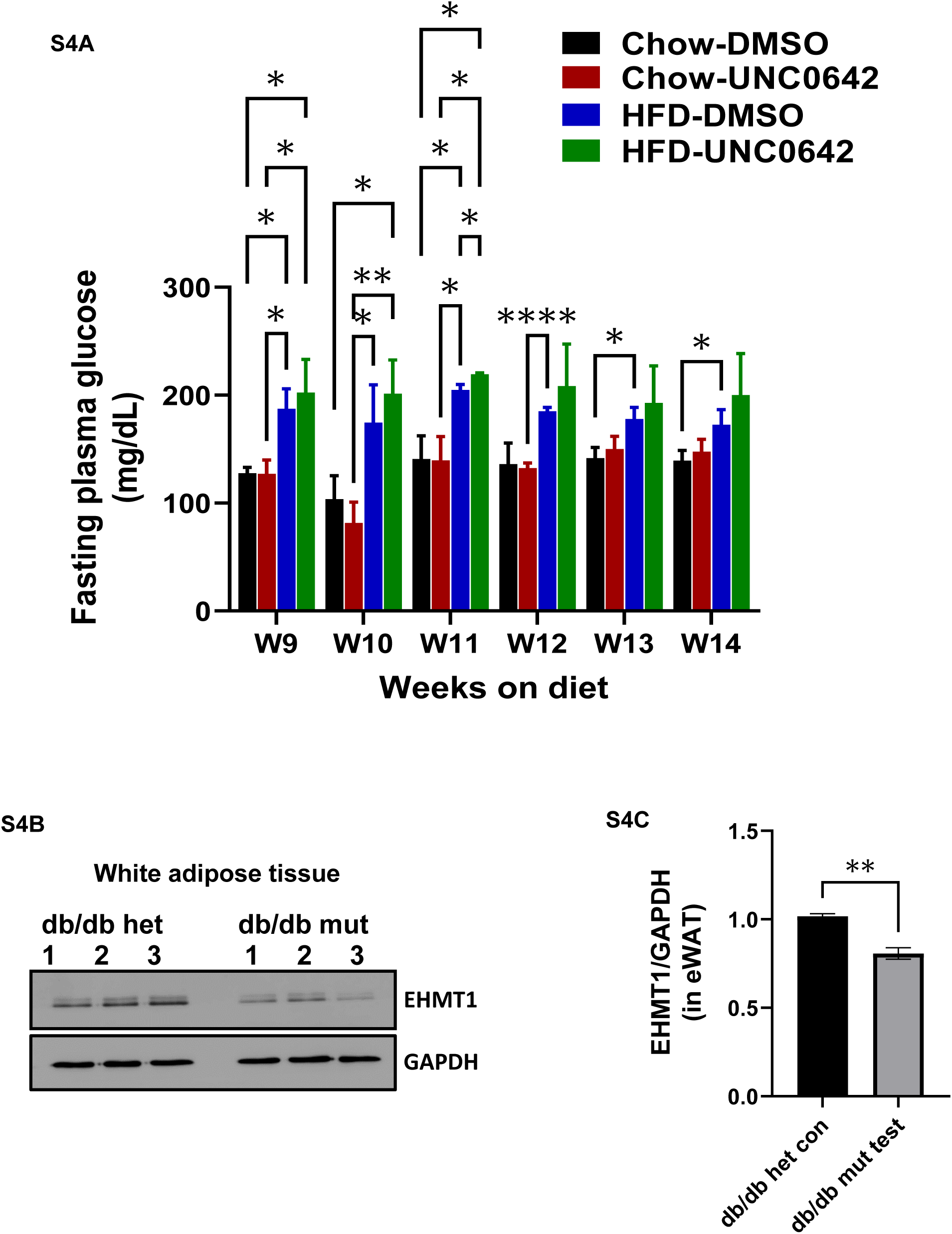
Supp Fig S4A: Fasting plasma glucose(mg/dl) from Chow fed-DMSO injected, Chow fed-UNC0642 injected, HFD fed-DMSO injected and HFD fed-UNC0642 injected mice (n=4 in each group). Supp Fig S4B: Western blot of 50pg total protein from white adipose tissue (epididymal white fat) of db/db het control and db/db mutant test mice (n=3) Supp Fig S4C: Densitometric analysis of EHMT1/GAPDH western band. Data are representative of minimum three independent experiments. Data are shown as Mean ± S.D. Unpaired t-test with Welch’s correction were performed wherever needed using Prsim 8.0 for statistical analyses, p ≤ 0.05 were marked with *, p ≤ 0.01 were marked with **, p ≤ 0.001 were marked with *** and considered significant.

## Supplementary Tables

Table 1. Mouse-specific primer sequences used for qPCR

Table 2. Primers used to amplify human EHMT1 promoter region carrying PPARƴbinding site and wild-type Sequence of Human pGL3-260 plasmid

Table 3. Primers used for performing site-directed mutagenesis in human pGL3-260 plasmid

Table 4. Sequences of respective substitution and deletion mutation regions in the H260-PGL3 plasmid

Table 5. Mouse-specific primer sequences used for CHIP qPCR analysis.

## Notes

### Competing Interest Statement

The authors have declared no competing interest.

